# Simple models of non-random mating and environmental transmission bias standard human genetics statistical methods

**DOI:** 10.1101/2024.10.16.618755

**Authors:** Richard Border, Jeremy Wang, Christa Caggiano, Sriram Sankararaman, Andrew J. Schork, Patrick Turley, Alexander S. Young, Daniel J. Benjamin, Andy W. Dahl, Noah Zaitlen

**Affiliations:** Department of Neurology, David Geffen School of Medicine, University of California, Los Angeles; Los Angeles, 90095, United States; Department of Computer Science, University of California, Los Angeles; Los Angeles, 90095, United States; Department of Computational Medicine, David Geffen School of Medicine, University of California, Los Angeles; Los Angeles, 90095, United States; Bioinformatics Interdepartmental Program, University of California, Los Angeles; Los Angeles, 90095, United States; Institute for Genomic Health, Icahn School of Medicine at Mount Sinai, New York, 10029, United States; Department of Human Genetics, University of California, Los Angeles; Los Angeles, 90095, United States; Institute of Biological Psychiatry, Mental Health Center Sct Hans, Copenhagen University Hospital–Mental Health Services CPH, 2100 Copenhagen, Denmark; Globe Institute, University of Copenhagen, 1350 Copenhagen, Denmark; Neurogenomics Division, The Translational Genomics Research Institute, Phoenix, 85004, United States; Department of Economics, University of Southern California; Los Angeles, 90089, United States; Center for Economic and Social Research, University of Southern California; Los Angeles, 90089, United States; Anderson School of Management, University of California Los Angeles; Los Angeles, 90095, United States; National Bureau of Economic Research; Cambridge, 02138, United States; Section of Genetic Medicine, Department of Medicine, University of Chicago; Chicago, 60637, United States

## Abstract

There is recognition among human complex-trait geneticists that not only are many common assumptions made for the sake of statistical tractability (e.g., random mating, independence of parent/offspring environments) unlikely to apply in many contexts, but that methods reliant on such assumptions can yield misleading results, even in large samples. Investigations of the consequences of violating these assumptions so far have focused on individual perturbations operating in isolation. Here, we analyze widely used estimators of genetic architectural parameters, including LD-score regression and both population-based and within-family GWAS, across a broad array of perturbations to classical assumptions, such as multivariate assortative mating and vertical transmission (parental effects on offspring phenotypes not mediated by genetic inheritance). We find that widely-used statistical approaches are unreliable across a broad range of perturbations, and that structural sources of confounding often operate synergistically to distort conclusions. For example, mild multivariate assortative mating and vertical transmission together can dramatically inflate heritability estimates and GWAS false positive rates. Further, GWAS will become progressively more polluted by off-target associations as sample sizes increase. Given these challenges, we introduce *xftsim*, a forward time simulation library capable of modeling a wide range of genetic architectures, mating regimes, and transmission dynamics, to facilitate the systematic comparison of existing approaches and the development of robust methods. Together, our findings illustrate the importance of comprehensive sensitivity analysis and present a valuable tool for future research.

## Introduction

Analysis of genome-wide data in large samples of unrelated individuals has become increasingly prevalent across the biomedical and social sciences, offering potentially valuable insights into the genetic bases of complex traits. The widespread adoption of these analyses has been propelled by the development of scalable statistical methods typically reliant on assumptions that make the challenging problem of relating individual differences in genetic markers with complex diseases and behaviors mathematically tractable. Examples of such assumptions have included purely additive genetic architectures, random mating, absence of indirect genetic effects, representative sampling, Mendelian randomization’s exclusion restriction assumption, and independence of genotypes and effects, among others. Understanding that the progress made possible by such simplifications must be balanced by sensitivity analysis and evaluation of alternative models, recent work has interrogated the consequences of perturbing such assumptions, including non-random mating (*1–5*), fine-scale population structure (*6*, *7*), participation bias (*8*, *9*), indirect genetic effects (*10–12*), and non-additivity (*13–18*).

Some of these assumptions appear to be relatively minor and not to distort our understanding of biology. For example, though dominance effects may be commonplace, methods assuming that common loci act additively on polygenic traits have been used to successfully find genetic pathways that are relevant to particular phenotypes (*14*, *18*). In other cases, realistic perturbations distort the conclusions we draw. For example, bivariate cross-trait assortative mating (xAM), which occurs when the process of mate selection results in cross-mate correlations among two phenotypes, can dramatically inflate genetic correlation estimates (*1*) and causes all loci causal for either phenotype to fail the exclusion-restriction criterion (*3*), a critical assumption for Mendelian-randomization-based causal inference.

Still, our understanding of the consequences of perturbing such assumptions is limited. First, only simple forms of such violations have been examined. For instance, despite evidence that assortative mating is widespread across the phenome, previous work has assumed that no more than two traits are involved in mate choice (*1*, *19*) or that phenotypes relevant to mate choice can be reduced to a linear dimension (*3*). Second, to the extent that such perturbations are examined, they tend to be examined in isolation, whereas multiple assumptions will be invalid for many traits of interest. For example, the highly-studied years of education phenotype is correlated across mates with a large number of traits, is thought to be subject to indirect genetic effects, and demonstrates complex causal relations with a number of other phenotypes—for example family wealth, which both impacts educational opportunities in childhood and is inherited through non-genetic mechanisms (i.e., bequests). As a result, despite the availability of increasingly large samples of unrelated genotyped individuals (*11*, *20*), the answers to longstanding questions such as whether or not years of education is affected by genes that influence BMI, remain unclear (*21*, *22*).

In the current manuscript, we analyze genome-wide association study (GWAS) effect estimates, as well as heritability and genetic correlation estimators, under simple perturbations of standard assumptions including phenotypes influenced by transmissible environmental factors (vertical transmission [VT], when parent phenotypes impact offspring phenotypes through the environment) and subject to multivariate cross-trait assortative mating (xAM, when mate choice is non-random with respect to multiple phenotypes). Toward this end, we develop *xftsim* (eXtensible Forward-Time Simulation framework), an open-source tool for scalable genotype / phenotype simulation in the context of complex architectures and mating / transmission dynamics (Table 1).

**Table 1.** Overview of xftsim library capabilities. Complete documentation is available online (23).

| **Feature** | **Approach** |
| --- | --- |
| Flexible mating regimes | Users may specify arbitrary target cross-mate cross-trait correlation structures across multiple phenotypes or phenotype components (e.g. additive-genetic or individual- / family-specific environmental factors); extensible via simulation state → mate pairing map abstraction |
| Vertical transmission / direct and indirect effects | Users may specify arbitrary causal relationships between phenotype components within and across generations; extensible via architectural component template class |
| Additive genetic architectures | Users may specify parametric infinitesimal and non-infinitesimal multivariate additive architectures or manually specify per-locus effects; extensible via additive-genetic architectural component template class |
| Non-additive genetic architectures | Users may specify quantitative gene-by-phenotype component (i.e. true polygenic index by component) interactions, including gene-by-gene or gene-by-environment effects; extensible via architectural component template class |
| Variable population size | Arbitrary distributions of numbers of mates- / numbers of offspring-per-individual; extensible via simulation state → realized matings abstraction |
| Realistic genotype data | Users may use seed simulations with real haplotype data or generate fully synthetic data and select between simple and map-based recombination schemes; *xftsim* provides utilities for reading and writing to plink binary and variant call format files |
| Population statistics / statistical estimators | *xftsim* includes functions for computing population statistics (e.g. true heritabilities, phenotypic / genetic correlations, cross-mate correlations, *et cetera*) and efficient implementations of widely-used statistical procedures, including GWAS, genetic (co)variance component estimation, and cross-validated polygenic index estimation; extensible via estimator template class |
| Sample ascertainment | *xftsim* enables users to specify arbitrary ascertainment regimes at the level of individuals or families for each estimator |

We show that, under very simple models of non-random mating and vertical transmission, widely used statistical genetic estimators (i.e., association tests, heritability estimators, and genetic correlation estimators) can generate misleading results and biased estimates, even when applied to nominally homogenous groups of unrelated individuals, and that increasing GWAS discovery sample size will only exacerbate such effects. In the process, we demonstrate that, for many phenotypes, xAM is not only multivariate (i.e., visible across many more than two phenotypes) but high-dimensional (i.e., cannot be described a single function of any set of observed or unobserved phenotypes); such higher-order xAM biases widely-utilized statistical procedures to a greater extent than bivariate xAM and accounts for a larger fraction of previously published estimates of genetic overlap among psychiatric disorders. Further, we demonstrate that perturbations of the aforementioned assumptions combine synergistically to further corrupt understanding of underlying biological relationships. For example, modest xAM (with cross-mate correlations of 0.2) and modest VT (accounting for 5% of phenotypic variance) together can result in genetic correlation estimates greater than 0.5 despite the complete absence of pleiotropy. Next, employing a heuristic example involving height, years of education, and wealth, we demonstrate that xAM and VT together can induce spurious genetic signals even for phenotypes with no causal genetic basis whatsoever. Finally, we explore the consequences of gene-environment interactions (G×E) and sample ascertainment in the context of nontrivial mating and transmission dynamics, as well as the extent to which family-based association studies are robust under such circumstances.

Together, our results highlight a blind spot in our current conceptualization of complex traits genetics: many existing methods are liable to give misleading results when applied to traits subject to complex social and behavioral dynamics. To address this, we provide a powerful simulation-based tool that enables researchers to assess the vulnerability of different methods to a comprehensive set of perturbations of classical assumptions affecting many human traits.

## Results

### xAM is multivariate

We began by quantifying the complexity of empirical cross-mate correlation structures across a limited collection of quantitative phenotypes chosen on the basis of previous evidence for xAM (*1*) and/or hypothesized relevance to selection (*24*). We applied canonical correlation analysis (CCA) to the cross-mate-cross-trait correlation structure for 34 phenotypes between 8,022 likely mate pairs with complete data in the UK Biobank (*25*) (Supplementary Table S1). The cross-mate correlation structure was high-dimensional: 22 canonical vectors evidenced significantly non-zero canonical correlations (*α*=0.05; see Supplementary Tables S2-S3 for complete results) and 8 canonical vectors were required to account for 90% of the sex-averaged relative canonical redundancy (the first canonical vector accounted for 36.7%; Figure 1a; see Supplementary Figures S1-S2 for supplemental analyses using a larger sample with imputed phenotype data). This implies that the full complexity of mate similarity, even across a limited number of phenotypes, cannot be fully characterized by one or two linear dimensions corresponding to physical appearance and/or a unidimensional social class construct as has been recently suggested (*26*). Instead, assortment is multifaceted, with many traits loading onto multiple canonical vectors. We further develop and differentiate the concepts of multivariate and multidimensional mating regimes in the Methods.

**Figure 1.**
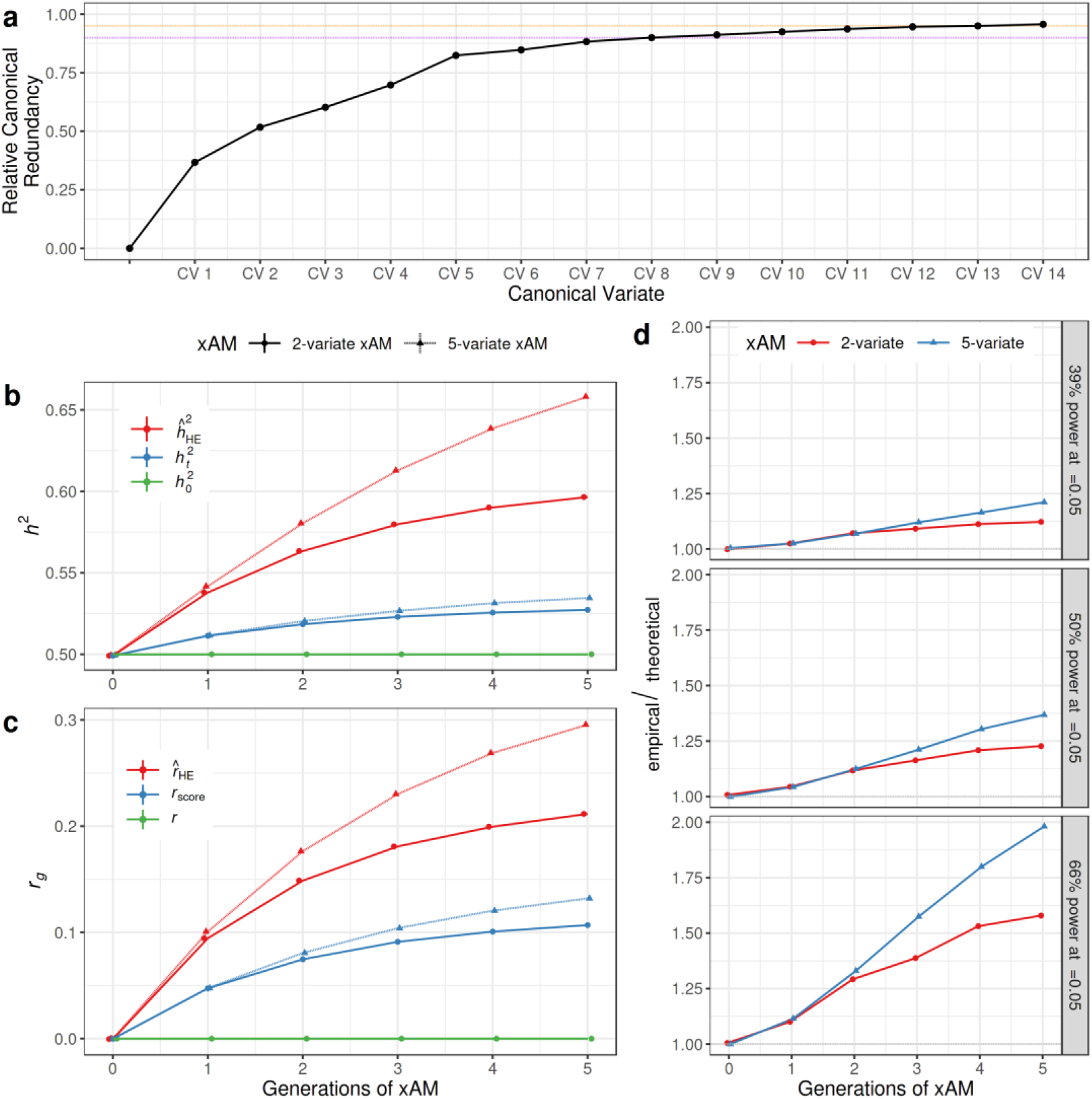
**(a)** Scree plot illustrating the sex-averaged canonical redundancy for each canonical vector relative to the total variance accounted for by all vectors. Eight (resp. fourteen) dimensions were required to explain 90% (resp. 95%) of shared variance across mates (purple [resp. orange] dotted lines). **(b)** True and estimated heritability over 0 to 5 generations of bivariate versus 5-variate xAM for simulated phenotypes with a true panmictic heritability (h^2^_0_) of 0.5, zero pleiotropy, and exchangeable cross-mate correlations of 0.2. As has been previously demonstrated, bivariate xAM increases the true generation-t heritability (h^2^), which is in turn overestimated by linear mixed model-based method-of-moments estimators such as Haseman-Elston and LD-score regression (h^^2^). However, relative to bivariate xAM, 5-variate xAM has a modestly larger effect on h^2^ but a substantially larger effect on h^^2^. **(c)** In the same simulation context, bivariate xAM increases the true correlation between polygenic scores (r_score_) and induces further upward bias in the corresponding method-of-moment estimate (r^_HE_), even when the true correlation between genetic effects (r_β_) is zero. As with heritability, 5-variate xAM has a modestly stronger effect on r_score_, the correlation between additive genetic factors, and induces substantially more severe upward bias in r^_HE_. **(d)** Ratio of empirical to expected type-I error rates for GWAS test statistics at Bonferroni significance across 0 – 5 generations of bivariate versus 5-variate xAM for simulated phenotypes. Panels correspond to the ratio of sample size to the number of causal variants-per-trait, with power increasing from top to bottom.

### Multivariate xAM induces severe biases relative to bivariate xAM

Having observed that multivariate xAM appears to be widespread across multiple phenotypic domains, we sought to quantify its potential impacts across a range of commonly applied methods for inferring genetic architecture by comparing bivariate xAM to 5-variate xAM. To enable this and all other simulations we present, we developed the eXtensible Forward-Time SIMulator (*xftsim*) software tool for human complex trait genetics research (*23*). *xftsim* allows users to model a wide variety of genetic architectures, mating regimes, and transmission dynamics as well as their effects on population statistics (e.g. allele-substitution effects, heritability, and genetic correlation) and estimators thereof. We provide an overview of the capabilities of *xftsim* in Table 1 and describe the implementation of multivariate mating and transmission regimes in the Methods.

Using *xftsim*, we simulated sets of two or five phenotypes subject to exchangeable cross-mate cross-trait correlations of 0.2 (i.e., such that all trait pairs are equicorrelated across mates), each with panmictic heritabilities *h*^2^_0_ = 0.5 and non-overlapping causal variant sets, partitioning the genome such that each variant was causal for one and only one phenotype. At each generation, we computed the true heritability *h*^2^ (the ratio of the additive genetic and phenotypic variance components) and true polygenic index (PGI) correlation *r*_score_ (i.e. the correlation between phenotypes’ additive genetic components), both of which increase over successive generations of assortment (*27*, *28*), as well as the corresponding Haseman-Elston estimators, *h^*^2^_HE_ and *r^*_HE_, which are biased upwards and yield results approximately equivalent to the widely used LD-score regression estimators (*2*, *29*).

Consistent with previous results (*1*), the true heritability increased across successive generations of bivariate xAM from *h*^2^_0_ =0.499 at generation zero to *h*^2^_0_ =0.527 at generation five. This increase was modestly larger under 5-variate xAM (*h*^2^_0_ =0.499 to *h*^2^_0_ =0.535; Figure 1b). On the other hand, though the estimated heritability was biased upwards under bivariate xAM (*h^*^2^_HE_=0.499 to *h^*^2^_HE_=0.597), the bias was substantially greater under 5-variate xAM (*h^*^2^_HE_=0.499 to *h^*^2^_HE_=0.658). We observed a similar pattern of results for genetic correlations, even when the true correlation between effects *r*_β_ was zero. First, five generations of bivariate xAM induced increases in the true polygenic index correlation (*r*_score_=0.107) and substantial upward bias in the estimated genetic correlation (*r^*_β_=0.212; Figure 1c). Again, the relative consequences under five generations of 5-variate xAM were moderate for the true polygenic index correlation (*r*_score_=0.107) but substantial for genetic correlation estimates (*r^*_β_=0.296). These results illustrate not only that the typical interpretation of genetic correlation estimates as measures of *r*_β_ (i.e., the extent to which loci similarly affect a pair of traits) is invalid for traits subject to assortment (*1*), but that this interpretation becomes increasingly tenuous when more traits are salient with respect to mate choice.

We also performed a series of simulations examining the impact of bivariate and 5-variate xAM on false-positive associations in the context of GWAS under variable statistical power (Figure 1d). Under the previously described simulation parameters, we performed association studies and computed false positive rates at a threshold of α = 0.05, varying statistical power from 39% to 66% by fixing sample size *n* and panmictic heritability while varying the average variance explained by each causal variant. By design, because each simulated variant was chosen to be causal for one and only one phenotype, most false positive associations for one trait will be “off-target” assocations with causal variants for another. We quantify excess false positives by examining the ratio of empirical to theoretical type I error rates (α^/α). Consistent with previous results, xAM induced false positive associations, and, as with the variance component analyses, 5-variate xAM generated stronger artifactual signals than bivariate xAM (α^/α=1.52 versus 1.32, respectively, averaging across power conditions). Critically, the false positive rate increases with statistical power: at 66% power, the type-1 error rate was 1.61 times larger than theoretical expectations under bivariate xAM and 1.98 times larger under 5-variate xAM. This implies that we should expect to detect larger numbers of spurious associations as sample sizes increase.

### Case study: genetic overlap among psychiatric disorders in the context of high-dimensional xAM

We next analyzed the potential consequences of high-dimensional xAM on six previously-studied psychiatric disorder groups with high cross-mate tetrachoric correlations (attention-deficit hyperactivity disorder [ADHD], alcohol use disorder [ALC], anxiety disorders [ANX], bipolar disorders [BIP], major depressive disorder [MDD], and schizophrenia [SCZ]; Supplementary Table S3) (*1*). Importantly, the empirical 6-variate cross-mate cross-correlation structure between disorders suggests that cross-mate similarity is widespread across pairs of disorders. We therefore compared expected genetic correlation estimates under xAM alone (*r*_xAM_; i.e., without pleiotropy) assuming bivariate versus 6-variate xAM (consistent with empirical cross-mate correlation estimates across the six disorders) and using empirical heritability estimates derived from twin and family studies (*1*). We also compared *r*_xAM_ to the empirical LD-score regression estimates of Grotzinger et al. (*r^*_empirical_) to determine the extent to which these estimates are consistent with xAM-induced artifacts alone (*30*) (Figure 2a). Finally, we characterized the impact of bivariate versus 6-variate xAM on correlations among GWAS effect estimates in the absence of pleiotropy.

**Figure 2.**
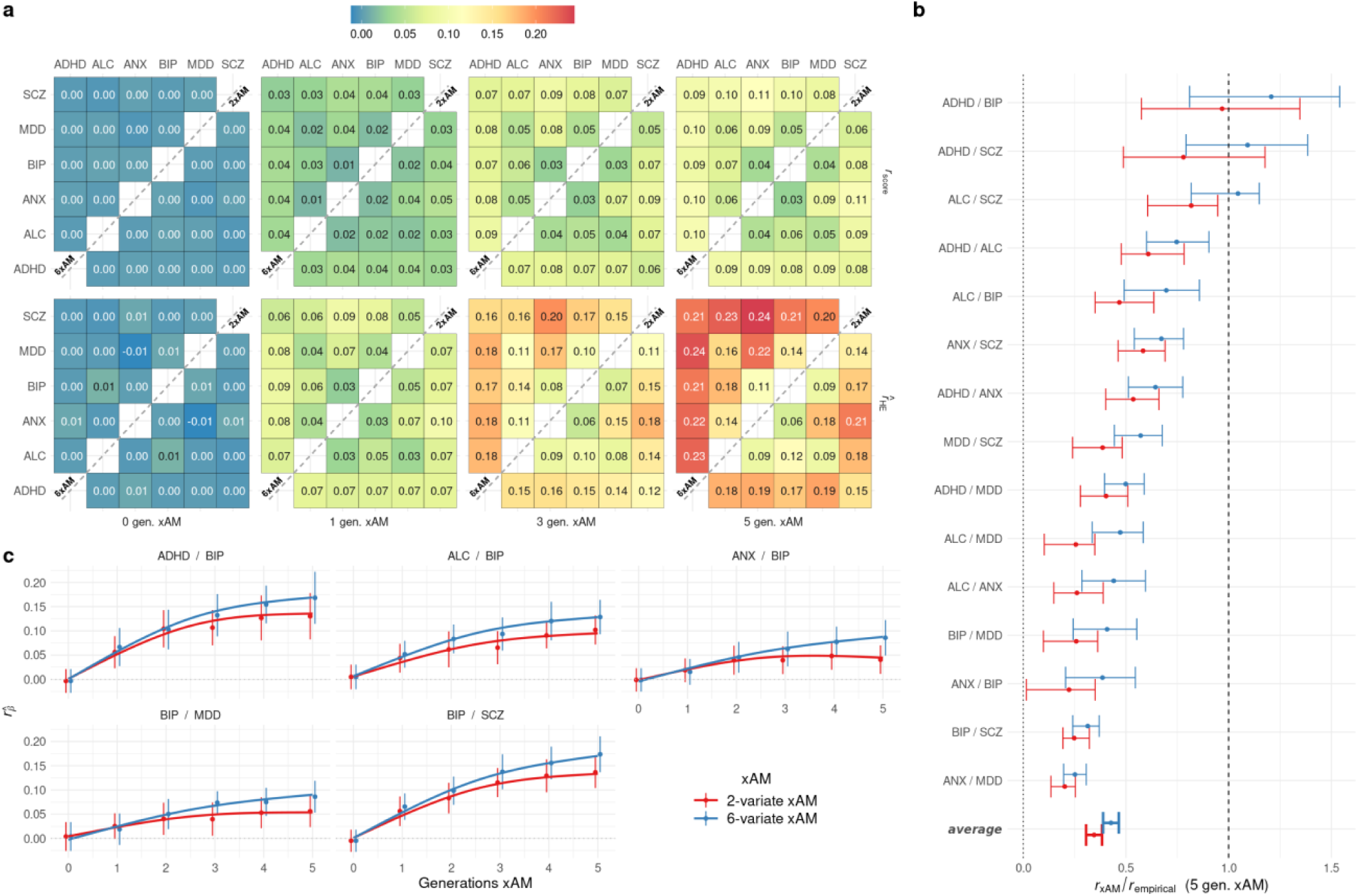
**(a)** True polygenic score correlations (r_score_; top row) and method-of-moments genetic correlation estimates (r^_HE_ ; bottom row) and among six psychiatric disorders after 0 – 5 generations of simulated xAM congruent with empirical spousal correlations. Superdiagonal elements reflect bivariate assortative mating (i.e. each cell corresponds to an independent set of simulations of xAM across the two referenced traits and no others) whereas subdiagonal elements reflect empirical 6-variate xAM (i.e. simulations of xAM across all six traits simultaneously). **(b)** Ratio of predicted genetic correlation under bivariate versus 6-variate xAM without pleiotropy (r_xAm_) to empirical LD-score regression genetic correlation estimates among psychiatric disorders (r_empirical_); values close to one reflect empirical estimates consistent with xAM alone, i.e. in the absence of pleiotropy. **(c)** Expected change in disorder prevalence under bivariate and 6-variate xAM. ADHD=attention-deficit hyperactivity disorder, ALC=alcohol use disorder, ANX=anxiety disorders, BIP=bipolar disorders, MDD=major depressive disorder, SCZ=schizophrenia.

As before, 6-variate xAM modestly increased correlations among true PGI when compared to bivariate xAM (e.g., for major depression [MDD] and anxiety disorders [ANX], *r*_score_=0.087 [*se*=1.58e-2] after five generations of 6-variate xAM versus *r*_score_=0.093 [*se*=1.54e-2] under bivariate xAM; Figure 2a). On the other hand, expected xAM-induced genetic correlation *estimates* were substantially higher under high-dimensional xAM (e.g., for MDD and ANX, *r*_xAM_=0.219 [*se* = 3.51e-2] after five generations of 6-variate xAM versus *r*_xAM_=0.176 [*se* = 4.29e-2] under bivariate xAM). Likewise, five generations of 6-variate xAM alone yielded larger genetic correlation estimates relative to those from the literature (*r*_xAM_/*r*_empirical_=0.427 [*se* = 2.32e-2] under 6-variate xAM versus *r*_xAM_/*r*_empirical_=0.345 [*se* = 2.36e-2] under bivariate xAM; Figure 2b). Notably, 6-variate xAM alone could fully account for empirical genetic correlation estimates for three trait pairs; for example, for the pair attention-deficit hyperactivity disorder and schizophrenia, five generations of 6-variate xAM yields estimates 1.09 [*se* = 0.228] times larger than previously published estimates (*31*)).

In addition to inducing false positives, xAM will induce directional bias in GWAS effect estimates across traits under assortment such that effect estimates become positively correlated. We illustrate this phenomenon for the six psychiatric traits in Figure 2c: although each of the six disorders was modeled as having unlinked, non-overlapping causal variants, five generations of bivariate xAM induced an average correlation among GWAS slope estimates of *r̅*_β^_=0.113 [*se* = 1.06e-2] across trait pairs. Again, this effect was more pronounced when modeling empirical xAM across all six traits: *r̅*_β^_=0.149 [*se* = 9.06e-3].

### Vertical transmission exacerbates xAM-induced biases

We next turn our attention to another mechanism underlying phenotypic variation that may also bias contemporary statistical genetic models: parents transmit more to their offspring than genetic material. We find that, in the presence of xAM, even weak parent-offspring (i.e., “vertical”) phenotype transmission (VT) dramatically compromises the interpretability of genetic estimators. We ran additional simulations examining the consequences of random mating (RM) versus 5-variate xAM (5xAM) in combination with multivariate VT explaining only 5% of the total phenotypic variance at generation 0 (Figure 3a; see Methods for detailed description of model parametrization). As with the 5-variate results presented in Figure 1b-1d, the five phenotypes were subject to exchangeable cross-mate cross-trait correlations of 0.2, each with panmictic heritabilities *h*^2^ = 0.5 and without pleiotropy across traits.

**Figure 3.**
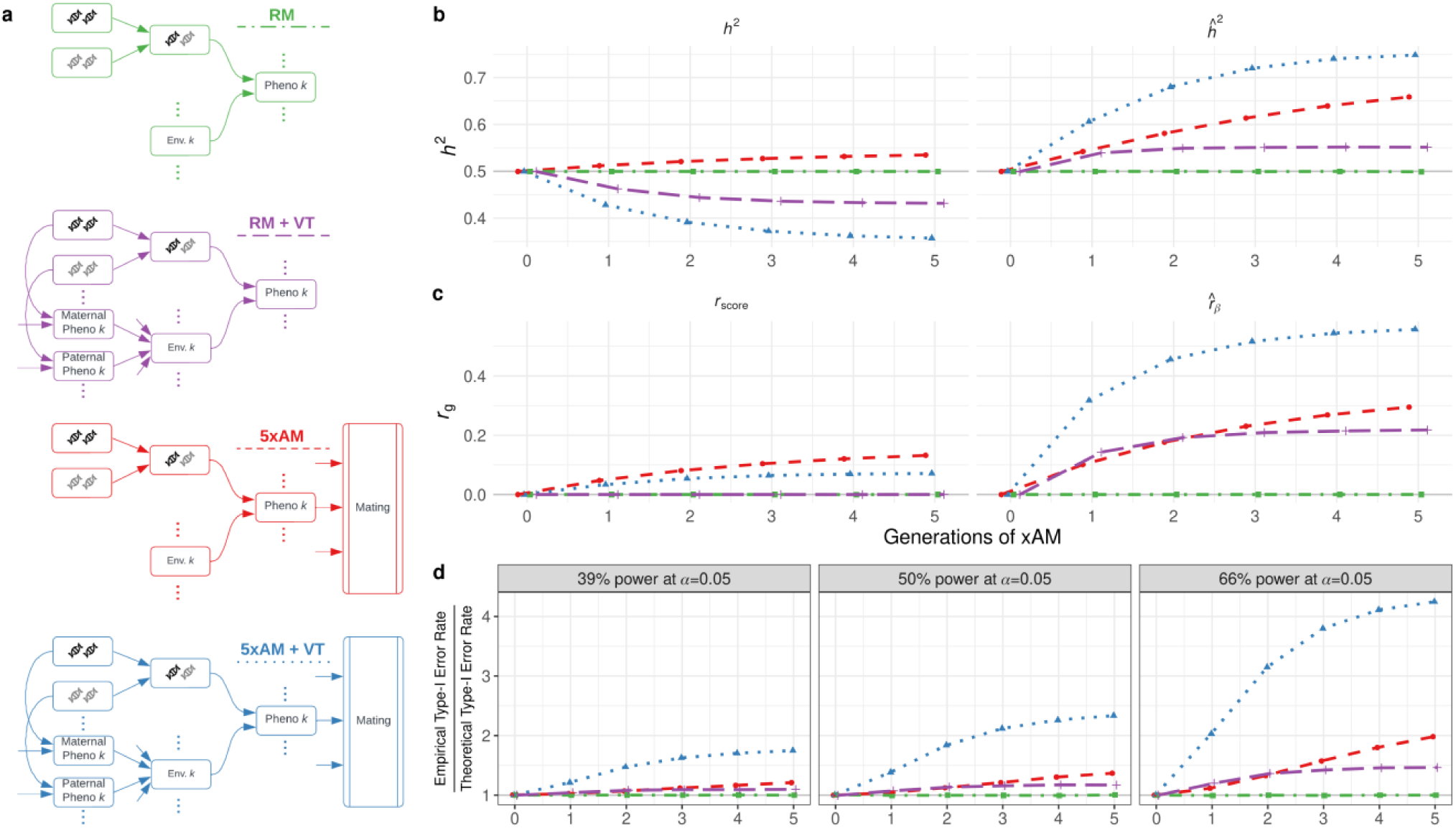
**(a)** Generative models for five phenotypes with orthogonal genetic effects. Phenotypes are subject to random versus 5-variate cross-trait assortative mating (RM versus 5xAM) and/or vertical transmission of environment from parent to offspring (VT). When present, VT accounts for 5% of the total phenotypic variance at generation zero **(b)** True (h^2^) and estimated (h^^2^_HE_) heritabilites over successive generations across generative models. **(c)**True polygenic score correlation (r_score_) and estimated effect correlation (r^_HE_) over successive generations across generative models. In all cases, the true effect correlation is exactly zero. **(d)** Relative inflation of empirical to nominal type-I error rates at nominal significance for GWAS with varying power (modulated here by the ratio of sample size to causal variants [n/m]). Note: The color of points in panels (b)-(d) correspond to those of the schematics in panel (a).

Compared to five generations of 5xAM alone, 5xAM + VT induced substantial countervailing differences in true and estimated quantities. After five generations of 5xAM + VT, whereas the true heritability fell from *h*^2^_0_ =0.500 to *h*^2^_0_ =0.396, the estimated heritability increased to *h^*^2^_HE_=0.749, with median upward bias of *h^*^2^_HE_ − *h*^2^_0_ =0.353 across simulation replicates, far larger than the already considerable bias induced by 5xAM of *h^*^2^_HE_ − *h*^2^_0_ =0.124. True heritability decreased because the variance explained by vertical transmission can increase more rapidly than the additive genetic variance under xAM. However, xAM combined with vertical transmission (which also induces indirect genetic effects when the parental phenotype is heritable, as it is here) can lead to strong gene-environment correlation that biases GWAS estimates and heritability estimators utilizing GWAS summary statistics (*5*, *32*). Likewise, 5xAM + VT decreased the true score correlations but increased the estimated effect correlations relative to 5xAM alone (*r*_score_=0.078 and *r^*_β_=0.513 versus *r*_score_=0.132 and *r*_score_=0.296 after five generations of 5xAM + VT versus 5xAM, respectively). When interpreted as an estimate of the correlation between direct genetic effects, this amounts to upward bias of *r^*_β_ − *r*_β_=0.513, but even when interpreted as an estimate of the true polygenic index correlation, this amounts to a bias of *r^*_β_ − *r*_score_=0.435.

We again performed a series of simulations examining GWAS false-positive rates under variable statistical power (Figure 3d). Here, as with the variance components analyses, the addition of VT on top of 5xAM induced substantially higher false positive rates relative to 5xAM alone (α^/α=2.779 versus α^/α=1.522, respectively, across power conditions). As before, the inflation of GWAS statistics was exacerbated in larger samples; for example, increasing power from 39% to 66% increased the relative false positive rate from α^/α=1.747 to α^/α=4.251 under 5xAM + VT.

Finally, we performed a number of supplemental simulations incorporating stronger VT (i.e., accounting for up to 20% of total phenotypic variance at generation zero) and/or linear gene-environment interactions (G×E; Supplemental Figures S3-S7; Supplemental Table S4; see Methods for details of generative model), a selection of which we highlight here. Although stronger VT (accounting for 20% of phenotypic variance), in combination with 5xAM, resulted in stronger inflation of genetic correlation estimates relative to weaker VT (accounting for 5% of phenotypic variance), the difference between estimates without VT and weaker VT was more substantial (*r^*_β_ −*r*_score_=0.600 under 5xAM + strong VT; *r^*_β_ − *r*_score_=0.435 under 5xAM + weak VT; *r^*_β_ − *r*_score_=0.164 under 5xAM alone). In other words, in the context of xAM, relatively weak and strong VT can have comparable effects, especially compared to zero VT. Additionally, incorporating G×E effects had a larger impact on heritability estimates than genetic correlation estimates, especially in the presence of xAM and VT (*h^*^2^_HE_ − *h*^2^_0_ =0.295 and *r^*_β_ −*r*_score_=0.624 versus *h^*^2^_HE_ − *h*^2^_0_ =0.403 and *r^*_β_ − *r*_score_=0.600 under 5xAM + G×E + strong VT versus 5xAM + strong VT, respectively).

### Case study: xAM and VT in a simplified model of years of education

Next, we illustrate how plausible multivariate mating patterns together with cross-generation phenotypic transmission complicates inferences in the context of a simplified, heuristic model of education years (EY), height, and wealth (Figure 4a). Using hypothetical simulation parameters, we center EY in light of the history of misattribution of social determinants of success to biological differences (*33*, *34*), as well as recent evidence that conventional population-based methods have overstated the extent to which EY is affected by genetic variants affecting other traits (e.g., BMI) (*11*, *21*, *22*). For simplicity, we assume that, in generation zero: one third of the variation in individual wealth is transmitted parental wealth and the remaining two thirds reflect individual-specific random variation; 1% of the variation in EY is additive genetic, two thirds is due to transmitted parental wealth and EY, and the remaining variation is individual-specific noise; and 60% of the variation in height is additive genetic with the remaining 40% individual-specific. Thus, the generation zero heritability of EY is exceedingly low (1%) and there are no genetic or other systematic influences on individual wealth apart from passed-down parental wealth. Height is 60% heritable at generation zero and exclusively interacts with EY and wealth through trivariate xAM (see Methods for further details).

**Figure 4.**
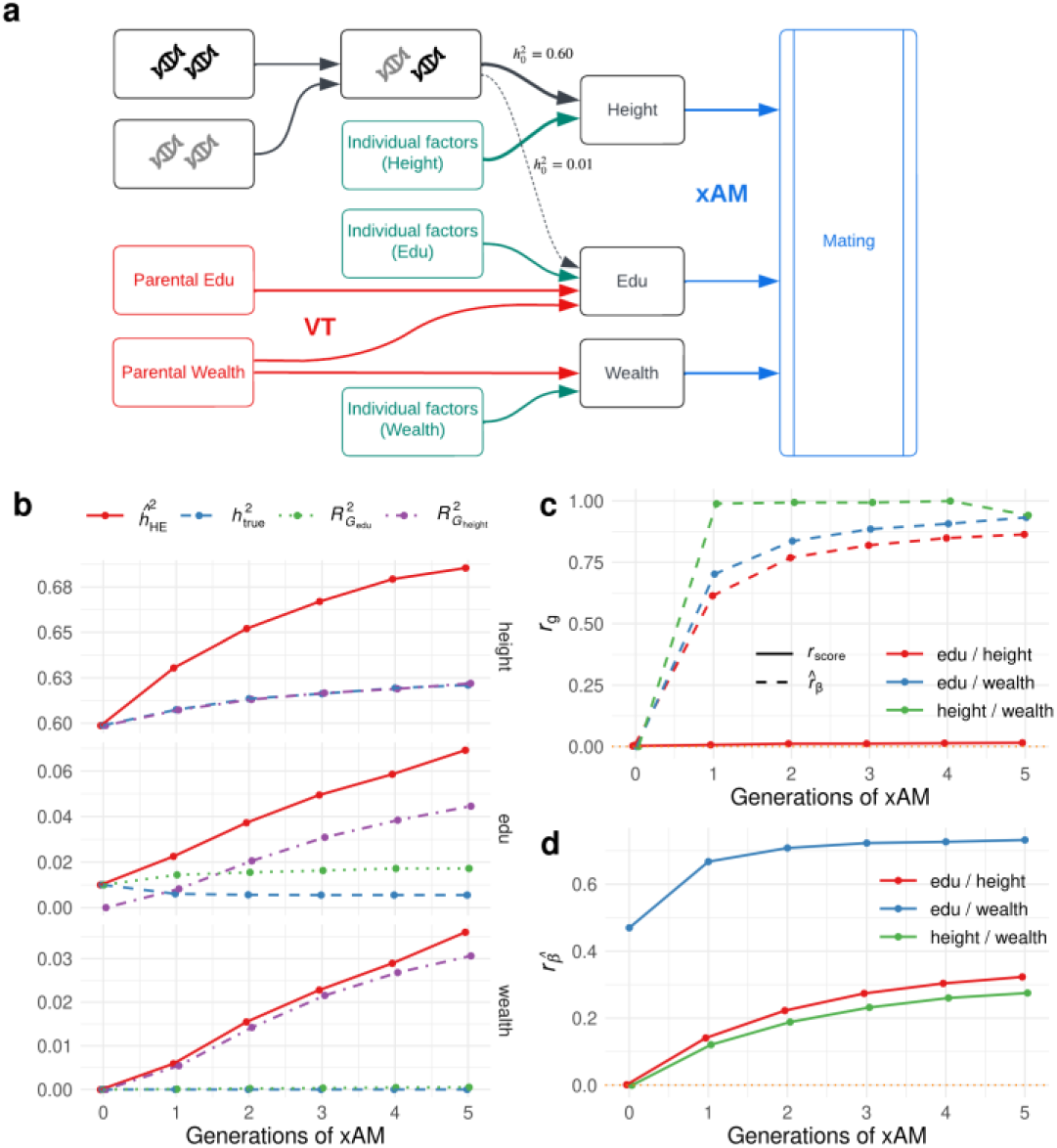
**(a)** Simplistic model of the joint architecture of wealth, education years (edu), and height. Height interacts with other phenotypes exclusively through cross-trait assortment (xAM), edu is trivially heritable (h^2^=0.01) but affected by parental education and wealth, and wealth has no genetic component but is transmitted from parents to offspring. **(b)** Estimated and narrow-sense heritability across successive generations of xAM, along with variance explained (R^2^) by true polygenic indices (PGI). **(c)** True PGI correlation and estimated genetic correlation across successive generations of xAM. Note that the true effect correlation between education and wealth in this case is identically one as wealth has no independent influences. **(d)** Correlation among GWAS effect estimates between height, edu, and wealth.

Across subsequent generations of xAM, the true heritability of height increased from *h*^2^_0_ =0.599 to *h*^2^_0_ =0.621. This increase was mirrored, but with upward bias, in heritability estimates, which increased from *h^*^2^_HE_=0.599 to *h^*^2^_HE_=0.686 (Figure 4b). On the other hand, the true heritability of EY decreased from *h*^2^_0_ =0.010 to *h*^2^_0_ =0.005, whereas the estimated heritability increased from *h^*^2^_HE_=0.010 to *h^*^2^_HE_=0.068. Likewise, estimated heritabilities for wealth, which had no genetic component whatsoever, increased from *h^*^2^_HE_=0.000 to *h^*^2^_HE_=0.036. For both traits, these estimates were larger even than the variance explained by regressing phenotype on the true polygenic indicess for either EY or height, which incorporates additive genetic variance and gene-environment covariance—e.g., after five generations, adjusted R^^2^=0.044 (resp. R^^2^=0.031) when regressing EY (resp. wealth) on the true height PGI. Genetic correlation estimates were also substantially biased upwards: though the true PGI correlation between height and EY increased from *r*_score_=0.0024 to *r*_score_=0.015 after five generations, the estimated genetic correlation increased from *r^*_β_=0.0053 to *r^*_β_=0.863 (Figure 4c). GWAS effect estimates also became correlated; for example, the correlation between effect estimates for height and wealth climbed from *r*_β^_=0.001 to *r*_β^_=0.273 (Figure 4d). In summary, our simplistic model demonstrates that phenotypic assortment and cross-generational transmission can lead to substantial biases in genetic estimates, even for traits with zero heritability (here, wealth).

### Within-sibship GWAS mitigates xAM and VT-induced biases with caveats

Given recent interest in within-family designs for addressing population-level confounds (*10–12*, *15*, *16*), as well as in the consequences of GWAS sample ascertainment (*8*, *9*), we performed simulations comparing population-based and within-sibship GWAS across the previously explored simple models, in both representative and ascertained samples. Our results suggest that, in contrast to population GWAS, within-sibship GWAS, under the simple models examined in the primary manuscript, are largely robust to some of the biases induced by xAM, VT, and/or G×E in representative samples: five generations of all combinations of these phenomena failed to increase association false positive rates, induce spurious correlations among slope estimates, or lead to substantial biases in slope effect estimates (Figures S7-S10). On the other hand, non-representative sampling substantially impacted both of these metrics in both population and within-sibship GWAS in a manner dependent on ascertainment regime, xAM, VT, and G×E (Figures S8-S10), though we emphasize that non-representative sampling, which we have only explored briefly here, appeared to have a substantially larger effect than the other perturbations.

Veller and colleagues recently demonstrated that sibling-difference GWAS estimate the local average treatment effect (LATE) of a given allele among individuals with at least one heterozygous parent at said locus, rather than the average treatment effect (ATE) in the population at large (*16*) (see also (*35*)). In the presence of G×E, then, sibling-difference GWAS estimates can differ from population average causal effects. Strictly speaking, these estimates are statistically unbiased, but with respect to the corresponding LATE; nevertheless, such estimates may be misleading when interpreted as population average causal effects or used for polygenic prediction. In supplementary analyses (see Methods), we show that, in the context of a model including VT and linear G×E, the relative difference between sibling-difference GWAS estimates and ATEs can be substantial at rare variants with large effects (Figures S11-S13). However, when causal variants have neutral effects on average and are symmetrically distributed with respect to allele frequency—as is the case for the simulations presented in the primary manuscript—the discrepancies between ATEs and sibling-difference GWAS estimates are modest and average out to zero across the genome (Figure S14). Under architectures where rare variants tend to have deleterious effects, sibling-difference GWAS derived PGI can differ substantially from PGI derived from true ATEs among carriers of rare variants of large effect (Figure S13). Of course, in the presence of true PGI-by-environment interactions, all polygenic predictors that fail to account for individuals’ environmental context can yield misleading predictions for individuals at environmental extremes (Figure S14), whether derived from within-family or population-based GWAS estimates (*36*).

## Discussion

The present manuscript demonstrates that perturbing common statistical genetic assumptions can cause widely-used estimators to yield misleading results, and that such effects can be compounded when multiple perturbations occur simultaneously. Further, empirical evidence suggests that multivariate xAM is widespread across numerous disease-relevant phenotypes, suggesting that many analyses will be affected by one or more such perturbations. Despite the broad array of scenarios examined here, we emphasize that the models we have considered, while less unrealistic than the standard models dominating the statistical genetics literature for many phenotypes, are at best crude approximations of the many social and biological factors that interact to shape genetic architecture and phenotypic diversity. With this in mind, we introduce the *xftsim* software package to enable comprehensive sensitivity analysis of existing methods and to facilitate the development of more robust approaches.

### Limitations of widely-used estimators under realistic complex trait architectures

Our findings underscore inherent limitations of current polygenic estimators, which largely treat genotypes, causal variant effects, and causal environmental factors as conditionally independent given genetic principal components and other covariates. On the other hand, for many disease-relevant phenotypes, such independence is unlikely to hold: assortative mating, which we and others have shown to be ubiquitous across a broad array of phenotypes (*1*, *37–39*) (see also Figure 1a, Supplementary Tables S2-S3), will induce sign-consistent long-range correlations between causal variants as a function of their effects (*1*, *2*, *19*, *27*, *40*, *41*), and the presence of indirect genetic effects (e.g., environmentally-mediated influences of parental genotype on offspring phenotype) will induce sign-consistent gene-environment correlations (*5*, *10*). While either of these dynamics in isolation can lead to misleading results, we have demonstrated that, together, even weak xAM combined with weak multivariate vertical transmission — which induces indirect genetic effects — can lead to dramatic overestimates of genetic correlation (Figure 3c) as well as dramatically inflate GWAS false positive rates (Figure 3d). These biases can impact all estimators that treat genotypes as fixed or independent of other parameters (i.e., causal variant and environmentally mediated effects). This does not imply that all inferences based on these approaches are erroneous; for example, five generations 6-variate xAM without pleiotropy yield genetic correlation estimates for major depression and anxiety disorders roughly 25% as large as empirical estimates, implying there is substantial shared genetic signal beyond that attributable to xAM (Figure 2a/b). However, our results do imply that many estimates will be substantially biased and often misleading. For example, for attention-deficit hyperactivity disorder and bipolar disorders, three generations of xAM congruent with empirical mating patterns (and zero pleiotropy) yielded estimates on par with empirical estimates (Figure 2a/b). For many behavioral phenotypes, conventional estimates may be more misleading than informative.

### Sample size is not enough

Since their inception, GWAS-focused ascertainment efforts have prioritized increasing sample sizes (*42*) because individual common variant effects are too small to be reliably detected until samples grow into the tens or hundreds of thousands. Moreover, large GWAS samples have enabled the field to progress beyond early candidate-gene research, which yielded numerous associations that later failed to replicate (*43–46*). Our results, which demonstrate that (i) multivariate xAM is widespread (Figure 1a/b) and (ii) xAM-induced false positives increase not only with dimension but with sample size (Figure 1e), complicate this narrative: if we are not there already, we will eventually enter a GWAS discovery regime in which increasing sample size will lead to more, not fewer, false positive associations. This is because we will identify causal variants, or variants in LD with causal variants, for secondary phenotypes under xAM with the focal trait.

### No method is robust across all scenarios examined

Assuming representative sampling and no G×E, within-family GWAS methods (e.g. regressing sibling phenotype differences on sibling genotype differences), appear relatively robust to the consequences of xAM in the context of the models examined here (i.e., do not identify off-target associations with other phenotypes under xAM jointly with the focal traits; Supplemental Figures S7-S8). At the same time, our results show that the within-family estimates can be affected by xAM and VT under some non-representative sampling regimes (Figures S8-S10), suggesting that further analysis is required to understand how such estimators behave in real samples. Further, our results build upon recent work on the interpretation of within-family GWAS estimates (*16*, *35*), illustrating that, in the presence of G×E, sibling-difference GWAS effect estimates for rare variants of large effect may differ substantially from population average effects (Supplemental Figures S11-S13), though we note that G×E can present challenges for all additive polygenic predictors (Supplemental Figure S14). We have developed *xftsim* to enable researchers to characterize and compare the performance of existing methods in the presence of complex mating and transmission dynamics and of facilitating the development of novel population-based and within-family methods able to accurately characterize complex trait architectures.

### Limitations and future directions

Limitations of our analysis provide avenues for future research. First, we have restricted our attention to analyses of nominally ancestrally-homogenous samples, likely obviating major sources of confounding across many disease and disease-relevant phenotypes. Moreover, though we examined several key sources of confounding, the parameter space of plausible perturbations to the classical additive genetic model is vast, and many other complexities could be explored. In future work, we intend to extend our analyses to include a broader array of phenotypic and genetic architectures, including epistasis, admixture, and spatially-varying effects on mating and environment. Further, we have limited our investigation to a subset of commonly used quantitative frameworks for deciphering the genetic basis of disease, leaving a number of questions open for future inquiry. For example, it remains unclear how global estimates of polygenicity are affected by perturbations to classical assumptions, as does the extent to which recently-developed fine-mapping methods that include an infinitesimal component are able to mitigate the confounding influences of multivariate xAM (*47*). Finally, there remains a need for robust scalable methods applicable to traits subject to the dynamics investigated in the present manuscript (and the many scenarios yet to be investigated); we have developed *xftsim* to facilitate these efforts. Despite these limitations, our work illustrates that understanding the genetic basis of complex traits will require the development of richer models accounting for the complexities the genome and the phenome, and that existing estimates should be interpreted carefully and critically.

### Software and code availability

The *xftsim* library is open-source under the GNU General Public License v3.0. *xftsim* is available on the Python Package Index (PyPI) (*48*) and at https://github.com/rborder/xftsim (*23*) and documented online at https://xftsim.readthedocs.io. Code necessary to reproduce our analyses and simulations is available at https://github.com/rborder/xftmanu_code_supplement (*49*). UK Biobank data were accessed under application 33127 and are available through the UK Biobank Access Management System at http://amsportal.ukbiobank.ac.uk.

## Acknowledgments

We would like to acknowledge Stephen Becker, Graham Coop, Sasha Gusev, Peter Hull, and Carl Veller for providing feedback and/or contributing to helpful discussions.

## Methods

### Analysis of empirical mating patterns

#### Software environment

Analyses were performed using Python v3.9.15 (*50*), Mathematica v13.3 (*51*), Hexaly Optimizer v12.5 (*52*), and R versions v4.1.2 or v4.2.0, depending on compute environments (*53*). Data cleaning and manipulation was performed using the *data.table* v1.14.8, *reshape2* v1.4.4, and *stringr* v1.5.1 R libraries and the *numpy* v1.22.4 and *pandas* v1.5.3 Python modules (*54–58*). Graphics were generated using the *ggplot2* v3.4.4 and *cowplot* v1.1.1 R libraries (*59*, *60*).

#### UK Biobank phenotypes

We selected 46 quantitative traits to examine across mates with previous evidence for assortment (*1*) and/or hypothesized to be under selection (*24*) available in the UK Biobank (*25*) (UKB; Supplementary Table S1), omitting phenotypes likely to be, or by definition, equivalent across mates (e.g., number of children and household income). Most phenotypes are straightforward to extract from corresponding UKB fields (see Supplementary Table S1) with the following exceptions. Blood and lipid phenotypes (HbA1C, glucose, triglycerides, and LDL cholesterol) were first adjusted for fasting time using a simple linear model. Triglycerides and LDL cholesterol were then further adjusted for statin use using literature-derived estimates (i.e., multiplied by 1.4286 and 1.1765, respectively; (*61–63*)). The definition of EY was complex and is provided in the Supplemental Code (*49*).

#### Mate identification in the UK Biobank

We adapted previously published approaches (*1*, *64*) to identify 39,710 pairs of individuals likely to be mates in the UKB. Briefly, we first selected sex-discordant pairs of unrelated individuals who reported living with their spouse and had the same values for distance to coast (field 24508), inverse distance to nearest road (field 24010), nearest distance to nearest major road (field 24012), and household size (field 709), and concordant on whether their property was rented versus owned (field 780).

#### Defining and quantifying high-dimensional xAM

It is not immediately clear when mate similarity might be termed “high-dimensional.” It is insufficient to require that many phenotypes are correlated across mates as this is expected even when a single construct is involved in assortment. For example, if mates sort on standing height, both standing and sitting height, which are highly correlated within individuals, will be correlated across mates as a result. We therefore operationalized mating regime dimensionality in terms of the number of independent linear dimensions required to account for the variance shared across mates. This allowed us to distinguish among multivariate mating regimes (i.e. those resulting in similarity across multiple traits): unidimensional regimes are well summarized by a single canonical variate, whereas multidimensional regimes require multiple canonical variates (Supplementary Figure S15). We assume linearity in each dimension, which numerical experiments suggest is likely to be conservative with respect to dimensionality—though there may be unidimensional nonlinear sorting regimes that cannot be summarized by a single canonical variate, such regimes are difficult to construct (Supplementary Figure S16).

To quantify xAM dimensionality in the UK Biobank, we applied canonical correlation analysis (CCA) across male and female mates and counted the sex-averaged number of canonical vectors required to explain 95% of the total variation in one mate’s phenotypes explainable by linear combinations of their partner’s phenotypes (by comparing the cumulative sum of the canonical redundancies to the total sum). We note that this approach is more conservative than simply counting the number of canonical correlations statistically different from zero; in a large sample, many canonical variates will be statistically significant; in this case 22 were significant at α=.05, though canonical variates 15 through 22 accounted for 3.69% of additional covariation beyond the 95.66% attributable to the first 14 canonical variates (Supplementary Table S2).

#### Cross-mate CCA procedure

To mitigate the impact of extreme outliers, all variables were Winsorized at the 0.005 and 0.995 quantiles. We then regressed out linear and quadratic effects of age for all phenotypes separately by sex to account for the fact that couples’ ages are correlated.

Beginning with 39,710 likely mate pairs in the UK Biobank, we then restricted our analyses to individuals with complete data across 34 phenotypes with missingness < 30%, resulting in 8,022 mate pairs without missing data. We performed CCA across mates using the *yacca* R library v1.4-2 (*65*), where the two matrices *X*, *Y* consisted of female and male mates’ respective measurements on the same phenotypes, with the exception of age at menarche, which was specific to females. We computed cumulative relative canonical redundancies as follows by dividing the vector of canonical redundancies (i.e., the vector where each element reflects the total variance in columns of *X* by columns of *Y* through each canonical variate, or vice-versa), by the total canonical redundancy (i.e., the total variance in columns of *X* by columns of *Y* through all canonical variates, or vice-versa), and averaging these vectors elementwise across males and females. The cumulative sums of the relative canonical redundancy vector thus indicate the fraction of total variance in one mate’s phenotypes explained by optimal linear combinations of their partner’s phenotypes. We assessed the shared dimensionality of the cross-mate phenotyping correlation structure by counting how many canonical variates were required to achieve a sex-averaged relative canonical redundancy of 95%. Canonical cross-loadings were analogously averaged across sexes.

We performed supplementary analyses in the full sample of 39,710 likely mate pairs after imputing missing values missing values separately by sex using predictive mean matching multiple imputation by chained equations (MICE) via the *mice* R package v3.16.0 (*66*). We performed ten imputation replicates separately for male and female mates, resulting in 100 cross-mate analyses; we present median estimates in Supplementary Figure S1. We also performed supplementary analyses based of five replicate imputations using nonlinear MICE via the *miceRanger* R package v1.5.0 (*67*), which yielded nearly identical results (Supplementary Figure S2).

### Simulating complex intergenerational dynamics and phenotypic / genetics architectures

#### The xftsim library

We developed the *xftsim* library for flexible and performant forward time simulation of arbitrarily complex mating patterns and trait architectures, including but not limited to: high dimensional xAM, direct and indirect genetic effects, transmissible environments, non-additive genetic effects, and annotation-dependent genetic architectures. We provide efficient implementations of several widely used polygenic estimators, including method of moments heritability and genetic correlation estimators, association studies, and cross-validated polygenic index prediction. The *xftsim* library is freely available and thoroughly documented online (*23*). We provide an overview of *xftsim*’s primary features in Table 1.

#### Simulating multivariate and multidimensional xAM

Some multivariate xAM regimes (i.e., unidimensional regimes; see Figure S14) can be modeled as sorting on a single linear combination of multiple phenotypes. In such cases, the problem of simulating xAM is computational simple: males and females may be independently ordered on a linear combination of the component phenotypes and random Gaussian noise and paired. We used this linear unidimensional approach for the simulations presented in Figure 1, Figure 3, and Supplemental Figures S3-S9.

While mathematically convenient, such regimes cannot reproduce realistically complex cross-mate correlation structures such as those observed empirically in the UKB (Figure 1a), across psychiatric disorders (Figure 2), or across the phenotypes presented in Figure 4. We formulate the problem of simulating high-dimensional xAM across an arbitrary number (*k*) of phenotypes as follows: Given *n* individuals’ phenotypes *Y* ∈ ℝ^*n*×*k*^, *n* potential mates’ phenotypes *Ỹ* ∈ ℝ^*n*×*k*^, and a target empirical *k* × *k* cross-mate cross-covariance matrix Ω^, we seek a permutation *P*^∗^ satisfying

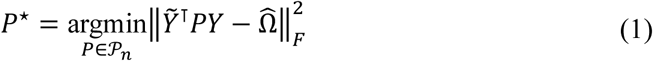

where *P*_*n*_ denotes the space of *n*-dimensional permutation matrices. In other words, we seek a pairing of individuals such that the resulting cross-mate cross-covariance matrix is close (in the least squares sense) to a (scaled) target cross-covariance matrix. Formulated this way, the problem of simulating arbitrary high-dimensional xAM is equivalent to the Quadratic Assignment Problem, a quadratic integer program well known in the fields of combinatorial optimization and operations research (*68*). Though finding the exact solution is NP-hard (*69*), we used the logistics optimization software Hexaly Optimizer v12.5 to find approximate solutions (*52*). In practice, we achieve cross-mate correlations matching target correlations to the third decimal place. Though the Hexaly Optimizer software is closed-source, academic licenses are available at no cost.

Both the linear unidimensional and quadratic integer program-based approaches to multivariate xAM are implemented in *xftsim*. Despite their implementation differences, both approaches yield nearly identical results when the target correlation structure is unidimensional: across 500 replicates-per-regime under the parameters described in the first row of Table 2, resulting cross-mate correlations, heritability, and genetic correlation estimates were indistinguishable across regimes (adjusted R^2^ when regressing outcomes on a mating regime indicator were all < 0.001 in magnitude).

**Table 2.** Phenotype-agnostic simulation model parameters. “2xAM”=bivariate cross-trait assortative mating, “5xAM”=5-variate cross-trait assortative mating, “RM”=random mating, “VT”= vertical transmission, “G×E”= gene-environment interaction, “num.”=number, “x-mate corr.”=cross-mate correlation, “var.”=variance. *See Figures 1c – 1e. ^†^See Figure 3. ^⁑^See Supplemental Figures S3-S5. ^‡^See Supplemental Figures S7-S10.

| Simulation | $K$<br>num. traits | $r$<br>x-mate | $\sigma_g^2$<br>additive | $\sigma_\varepsilon^2$<br>noise | $\theta$<br>VT var. | $\phi$<br>G×E var. |
| --- | --- | --- | --- | --- | --- | --- |
| 2xAM* | 2 | 0.2 | 50% | 50% | 0% | 0% |
| 5xAM** | 5 | 0.2 | 50% | 50% | 0% | 0% |
| RM† | 5 | 0.0 | 50% | 45% | 5% | 0% |
| RM + VT† | 5 | 0.0 | 50% | 45% | 5% | 0% |
| 5xAM + VT†* | 5 | 0.2 | 50% | 45% | 5% | 0% |
| 5xAM + G×E†* | 5 | 0.2 | 50% | 45% | 0% | 5% |
| 5xAM + VT + G×E†* | 5 | 0.2 | 50% | 40% | 5% | 5% |
| 5xAM + VT (20%)* | 5 | 0.2 | 50% | 30% | 20% | 0% |
| 5xAM + G×E (20%)* | 5 | 0.2 | 50% | 30% | 0% | 20% |
| 5xAM + VT (5%) + G×E (20%)* | 5 | 0.2 | 50% | 25% | 5% | 20% |
| 5xAM + VT (20%) + G×E (5%)* | 5 | 0.2 | 50% | 25% | 20% | 5% |
| 5xAM + VT (20%) + G×E (20%)* | 5 | 0.2 | 50% | 10% | 20% | 20% |

#### Phenotype-agnostic simulation parametrizations

For a given offspring phenotype vectors *Y*_*k*_ denote the corresponding maternal and paternal phenotype vectors by *Y*^∗^, *Y*^∗∗^, respectively. For the results presented in Figures 1 and 3 and Supplemental Figures S3-S9, the phenotypic generative model for *K* traits within each generation was of the form

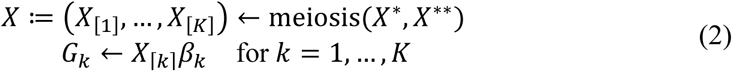

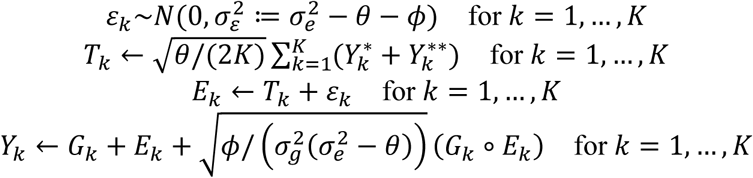

where each vector of causal effects β_*k*_ for the *k*th independent set of *m* standardized SNPs *X*_[*k*]_ is distributed β_*k*_∼*MVN*(0_*m*_, σ_g_^2^/*m*). At generation zero the variance components represent: σ_e_^2^, the total non-additive genetic variance; σ^2^, the individual-specific noise variance; σ_g_^2^, the additive genetic variance; θ, the variance attributable to the combined effects of transmitted parental effects; and Φ, the variance attributable to gene-environment interaction. Given females’ phenotypes *Y̅*_1_, …, *Y̅*_*K*_ and males’ phenotypes *Y̿*_1_, …, *Y̿*_*K*_, individuals were paired such that the cross-mate phenotypic cross correlation cor(*Y̅*_*k*_, *Y̿*_*l*_) was identically equal to *r* across all pairs of phenotypes *k*, *l* = 1, …, *K*. Across all simulations the total additive genetic and non-additive-genetic variances components were fixed to σ_g_^2^ = σ_e_^2^ = 1; full parameter specifications are detailed in Table 2. Statistical estimates reported were averaged across individual phenotypes for univariate quantities (e.g., true and estimated heritabilities and GWAS false positive inflation indices) and across all distinct pairs of phenotypes for bivariate quantities (e.g., true polygenic index correlations, estimated genetic correlations, and GWAS slope estimate correlations) to minimize sampling error.

#### Psychiatric phenotype simulation parameters

The psychiatric disorder simulations presented in Figure 2 assume a simple genetic architecture where each disorder was comprised of independent additive genetic and non-genetic variance components and each additive component reflected product of *m* unique (i.e., phenotype-specific, ensuring zero pleiotropy) standardized SNPs with Gaussian effects. Simulations were parametrized using previously published empirical family-based heritability estimates and cross-mate cross-correlation tetrachoric correlation estimates (*1*), and mating was simulated using the quadratic integer programming approach described above. Statistical estimators were applied directly to continuous liabilities (as opposed to dichotomized phenotypes) to minimize variance.

#### Height, education, and wealth simulation parameters

Using subscripts to denote height, education, and wealth by *Y*_h_, *Y*_e_, *Y*_w_, respectively, and again distinguishing mates by superscript asterisks, the simulations presented Figure 4 assumed the generative model

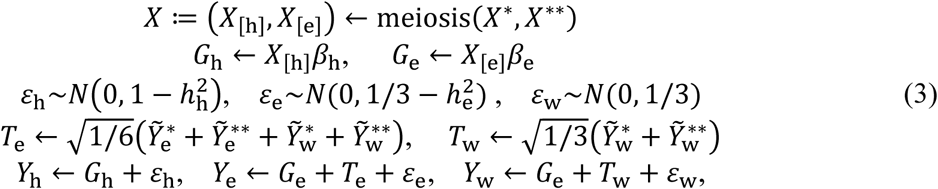

where the heritabilities of height and education were set to *h*^2^_0_ =0.6, *h*^2^_0_ =0.01, and the remaining parameters were chosen such that, at generation zero, 0.67 (resp., 0.32) of the variance in education was attributable to vertically-transmitted education and wealth (resp. independent noise), and 0.67 (resp. 0.33) of the variance in wealth was attributable to transmitted wealth (resp. independent noise). Tildes denote standardized quantities. Mating was simulated using the quadratic integer programming approach described above to achieve the cross-mate cross-correlation structure

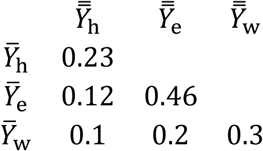

where the correlations across height and education were based on our empirical estimates in the UKB. In the absence of empirical estimates for wealth (only household-level metrics are available in the UKB, which are identical across mates by construction), plausible values were substituted. We emphasize that this analysis is intended to characterize the behavior of statistical estimators under nontrivial models incorporating xAM and VT, not to provide an empirical analysis of the height, wealth, and EY. We further remark that even a simplistic approach such as the above will be less unrealistic than the additive genetic random mating models which dominate the statistical genetics literature.

### Discrepancies between population average effects and within-sibship GWAS estimates induced by vertical transmission and G×E interactions

Veller and colleagues (*16*) recently demonstrated that sibling-difference GWAS estimates the local average treatment effect (LATE) of a given variant among individuals with at least one parent heterozygous at said locus. Intuitively, this is because siblings with homozygous parents will always (in the absence of mutation) have the same genotypes. The LATE need not agree with the average treatment effect (ATE), the causal effect of the variant in the general population (corresponding to the hypothetical experiment where an embryo’s genome is altered at the specific locus via gene editing). Particularly, in the presence of G×E effects, the LATE and ATE can differ when the average environment among individuals with heterozygous parents differs from the average in the general population. Such discrepancies between LATEs and ATEs can create challenges for clinical applications of within-family GWAS-derived PGI, where we want predictions that generalize across patients regardless of parental genotype.

One important scenario in which this is likely to occur is when allele frequencies vary across population strata with differing environments, as is the case when socioeconomic opportunities and resources are unevenly distributed with respect to genetic ancestry. This is a crucial topic for further research. Another scenario, which has particular relevance to the present manuscript, is vertical transmission of a heritable phenotype, which can induce correlations between parent genotype and offspring rearing environments. However, in the simulations incorporating vertical transmission presented in Supplemental Figure S6, we did not detect substantial differences between true ATEs and sibling-difference GWAS estimates. Here, we show that substantial discrepancies are expected under alternative genetic architectures, focusing on two illustrative examples.

#### Individual variants accounting for substantial phenotypic variation

Consider the following model for one generation of vertical transmission in the presence of additive and G×E effects:

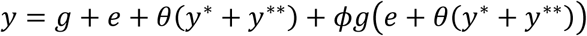

Here *y*^∗^, *y*^∗∗^ denote parents’ phenotypes and *g* and *e* are independent zero-mean Gaussians. We assume G×E (controlled *by* Φ) but not vertical transmission (θ) was present in the previous generation such that

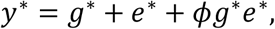

and likewise for *y*^∗∗^. We note that this model is the univariate version of the model described in Figure 3. We define the following variance components sequentially:

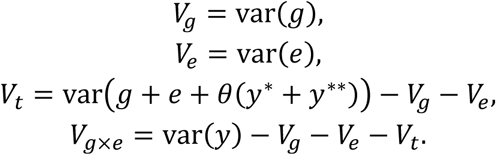

Decompose *g* into *g* = *g*_*j*_ + *g*_−*j*_ such that *g*_*j*_ : = *X*_*j*_β_*j*_ (a variant of large effect) is Binomial and *g*_−*j*_ : = ∑_*i*≠*j*_ *X*_*i*_ β_*i*_ (the additive genetic component due to all other variants) is Gaussian, and assume these variables are independent. Parametrize the signed fraction of additive genetic variance due to *g*_*j*_ by λ_*j*_ : = sgn (β_*j*_)var(*g*_*j*_)/*V*_*g*_and denote the allele frequency of *X*_*j*_ by *f*_*j*_. Using the laws of Mendelian inheritance and doing some nasty algebra reveals that there are four values of Φ, θ that satisfy given values of *V*_*g*_, *V*_e_, *V*_*t*_, *V*_*g*×e_, λ_*j*_, and *f*_*j*_, counting multiplicities (see Supplemental Code for a Mathematica notebook deriving these results (*49*)).

Under this model, we obtain yield unique ATE and LATE values:

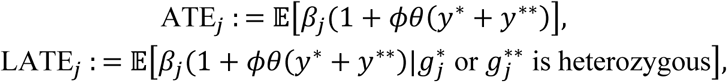

noting that sibling-difference GWAS estimates the later. Figures S11-S12 display the relative difference of the these quantities (LATE_*j*_ − ATE_*j*_)/|ATE_*j*_| as a function of λ_*j*_, the signed fraction of genetic variance due a given variant, for fixed *V*_*g*_ = 0.5, *V*_e_ = 0.3, *V*_*t*_ = 0.1, *V*_*g*×e_ = 0.1 and varying MAF (only *minor* allele frequency matters here, the results are symmetric in *f*_*j*_ about 0.5). All together, we note that:

1. The relative bias (i.e., the expected relative error in a sibling-difference GWAS if interpreted as a measure of the ATE) is not uniquely determined by the variance components / effect / MAF alone. This makes sense as we have a dynamic model where variance components change over time.
2. The effects can be substantial for rare variants and/or variants that account for a large fraction of additive variance and are small for common variants with small effects (and approach zero as allele frequency approaches 0.5).
3. Fixing all other variables, for any given solution, the relative bias appears to be monotone in λ_*j*_ in a neighborhood of the origin and crosses the origin for any given solution. Further, the sign of the relative bias is a function of the sign of the causal effect.
4. This kind of model does not appear to induce systematic bias in common variant effect estimates derived from sibling-difference GWAS as the half the effects will be upward-versus downward-biased.
5. On the other hand, the bias for a single variant of large effect will be greater than the sum of that of many variants of smaller effects (in the same direction) accounting for the same amount of phenotypic variance in aggregate.

We use the term “bias” loosely here, assuming that sibling-difference GWAS slopes are interpreted as ATE estimates. Strictly speaking, these estimators yield unbiased estimates of the LATE in individuals with at least one heterozygous parent.

Note that because PGI sum across variants with upward and downward bias, sibling-difference GWAS will yield unbiased estimates of the “true PGI” for highly polygenic traits without large effect variants (and in the absence of population structure or other confounders), if we define the true PGI to be the sum of genotypes multiplied by the corresponding ATEs. However, in the presence of substantial *G*×*E* effects, this population average PGI can be extremely misleading for individuals at environmental extremes, which we demonstrate in Figure S14 for Φ = 0, θ = 2^−1/2^. This effect is not specific to sibling-difference GWAS and can compromise any polygenic predictor that models genetic liability independent of environmental context.

#### Rare variants with large unstandardized effects

We have so far demonstrated that, when additive genetic effects have zero mean and are symmetrically distributed with respect to allele frequency, discrepancies between ATEs and the LATEs estimated by sibling-difference GWAS average out and sibling-difference GWAS-derived PGI will yield unbiased estimates of the ATE-based PGI (Figure S14). However, when rare alleles tend to have deleterious effects, these discrepancies need not cancel out. To demonstrate one such scenario, we simulated 100 causal variants with allele frequencies following a truncated beta distribution with shape parameters 0.4 and 1.0 on the interval [0.001, 0.98], with Gaussian standardized effects with mean −0.20 and variance *V*_*g*_ = 0.5 under the previously described model of vertical transmission and *G*×*E* with *V*_e_ = 0.3, Φ = 0.32, and θ = −0.70, yielding *V*_*t*_ = *V*_*g*×e_ = 0.1. We note that there are many potential causal effect distributions and that we selected this one simply for the sake of generating a clear example. Under these circumstances, despite accounting for little phenotypic variation in the population, the discrepancies between ATEs and sibling-difference GWAS effects for rare variants with large (unstandardized) effects will be negative on average, leading to downward-biased estimates of ATE PGI among carriers (Figure S13).

## Supplementary Tables

**Table S1.** Phenotype definitions. Across all phenotypes examined, we relied on a number of procedures to account for repeated measurements and/or missing data in the UK Biobank. We denote UK Biobank fields using the f. <FIELD>. <INSTANCE>. <INDEX>identifiers provided with the data. For example, f.709.0.0 through f.709.3.0 indicates each of the four possible times participants were asked about household size. Depending on the field, we either used the earliest available measurement, which we denote f. <FIELD>.E. <INDEX>, or the average measurements, which we denote f. <FIELD>.A. <INDEX>. We omit the index number when only one item is available, as is the case for many of the phenotypes, and use an asterisk when responses are array-valued.

**Table S2.** Cross-mate canonical correlation analysis results. Rows correspond to canonical variates. Columns include sex-specific canonical variate adequacies, canonical redundancies, and phenotype loadings (i.e., structural correlations).

**Table S3.** Cross-mate correlation estimates. For each trait pair, we provide sex-specific within individual and cross-mate Pearson and Spearman correlation estimates and corresponding standard errors.

**Table S4.** Median true and estimated variance components before and after five generations under random mating (RM), bivariate cross-trait assortative mating (2xAM), or 5-variate xAM (5xAM), with or without multivariate vertical transmission (VT) and/or gene-environment interactions (G×E) accounting for 5% or 20% percent of phenotypic variation at generation zero. When present, xAM is such that the cross-mate cross-correlations are 0.20 across all phenotype pairs. All phenotypes have additive genetic heritabilities of 0.50 at generation zero and orthogonal genetic effects.

| Scenario | Generation | $h^2$ | $\hat{h}^2$ | $\hat{h}^2 - h^2$ | $r_{\text{score}}$ | $\hat{r}_\beta$ | $\hat{r}_\beta - r_{\text{score}}$ |
| --- | --- | --- | --- | --- | --- | --- | --- |
| RM | 0 | 0.500 | 0.500 | 0.000 | 0.000 | 0.000 | 0.000 |
|  | 5 | 0.500 | 0.500 | 0.000 | 0.000 | 0.000 | -0.001 |
| RM + VT (5%) | 0 | 0.500 | 0.500 | 0.000 | 0.000 | 0.000 | 0.000 |
|  | 5 | 0.442 | 0.552 | 0.110 | 0.000 | 0.199 | 0.199 |
| 2xAM | 0 | 0.499 | 0.499 | 0.000 | 0.000 | 0.000 | 0.000 |
|  | 5 | 0.527 | 0.597 | 0.069 | 0.107 | 0.212 | 0.105 |
| 5xAM | 0 | 0.500 | 0.500 | 0.000 | 0.000 | 0.000 | 0.000 |
|  | 5 | 0.535 | 0.658 | 0.124 | 0.132 | 0.296 | 0.164 |
| 5xAM + G×E (5%) | 0 | 0.502 | 0.502 | 0.000 | 0.000 | 0.000 | 0.000 |
|  | 5 | 0.534 | 0.658 | 0.124 | 0.133 | 0.297 | 0.165 |
| 5xAM + G×E (20%) | 0 | 0.543 | 0.543 | 0.000 | 0.000 | 0.000 | 0.000 |
|  | 5 | 0.570 | 0.714 | 0.145 | 0.141 | 0.317 | 0.175 |
| 5xAM + VT (5%) | 0 | 0.500 | 0.500 | 0.000 | 0.000 | 0.000 | 0.000 |
|  | 5 | 0.396 | 0.749 | 0.353 | 0.078 | 0.513 | 0.435 |
| 5xAM + VT (5%) + G×E (5%) | 0 | 0.499 | 0.500 | 0.000 | 0.000 | 0.000 | 0.000 |
|  | 5 | 0.371 | 0.723 | 0.352 | 0.073 | 0.525 | 0.452 |
| 5xAM + VT (5%) + G×E (20%) | 0 | 0.535 | 0.535 | 0.000 | 0.000 | 0.000 | 0.000 |
|  | 5 | 0.297 | 0.627 | 0.330 | 0.060 | 0.555 | 0.495 |
| 5xAM + VT (20%) | 0 | 0.500 | 0.499 | 0.000 | 0.000 | 0.000 | 0.000 |
|  | 5 | 0.247 | 0.650 | 0.403 | 0.034 | 0.634 | 0.600 |
| 5xAM + VT (20%) + G×E (5%) | 0 | 0.488 | 0.488 | 0.000 | 0.000 | 0.000 | 0.000 |
|  | 5 | 0.164 | 0.459 | 0.295 | 0.029 | 0.653 | 0.624 |
| 5xAM + VT (20%) + G×E (20%) | 0 | 0.500 | 0.500 | 0.000 | 0.000 | 0.000 | 0.000 |
|  | 5 | 0.064 | 0.195 | 0.130 | 0.020 | 0.677 | 0.658 |

**Figure S1.**
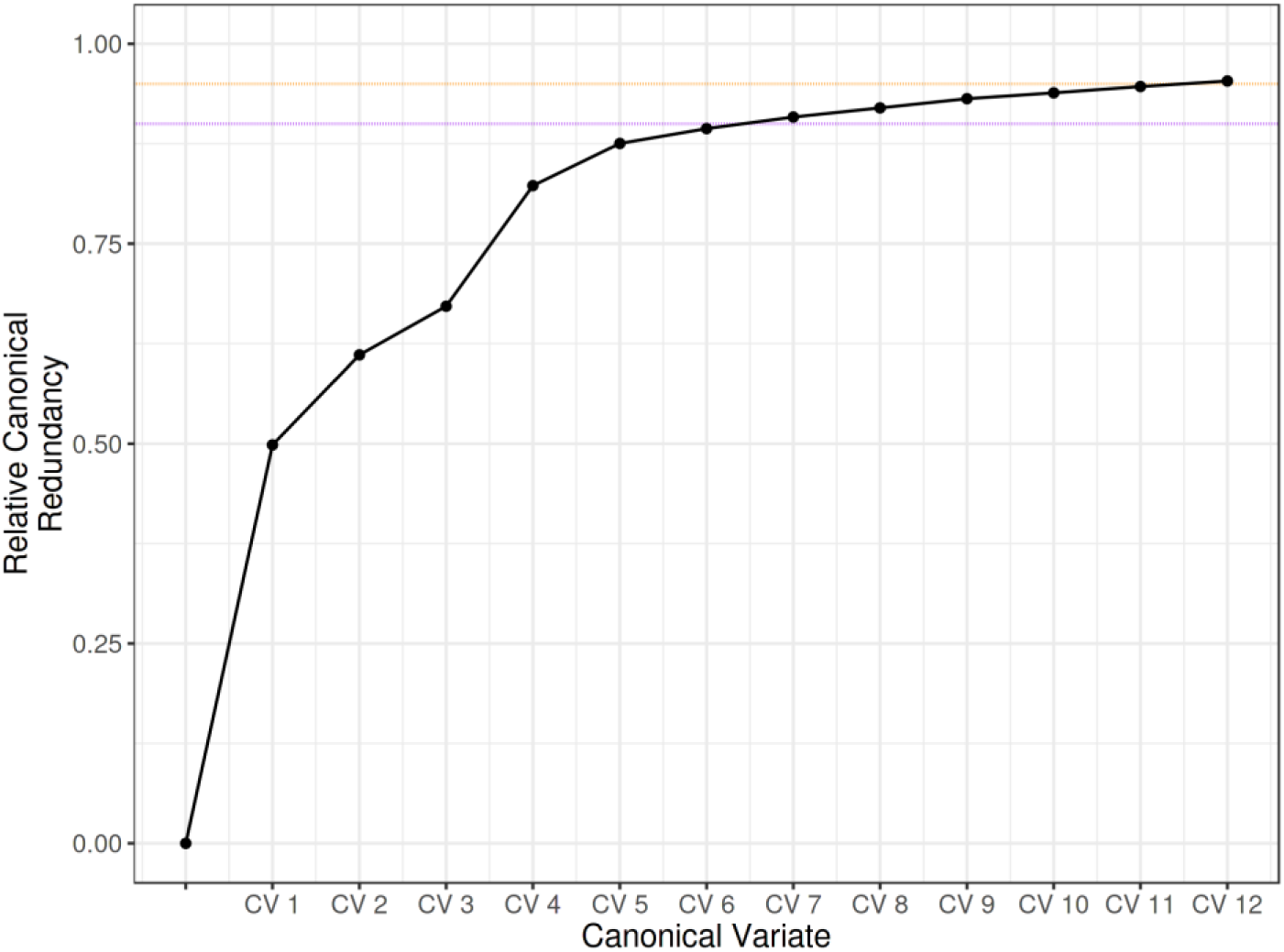
Cross-mate CCA scree plot based on multiply imputed data using predictive mean matching multiple imputation by chained equations. Vertical axis reflects sex-averaged cumulative relative canonical redundancy. Twelve linearly independent canonical variates are required to account for 95% of the variance explicable in one mate’s phenotype by linear combinations of their partner’s phenotypes. Values reflect median redundancies across 5 × 5 imputed datasets.

**Figure S2.**
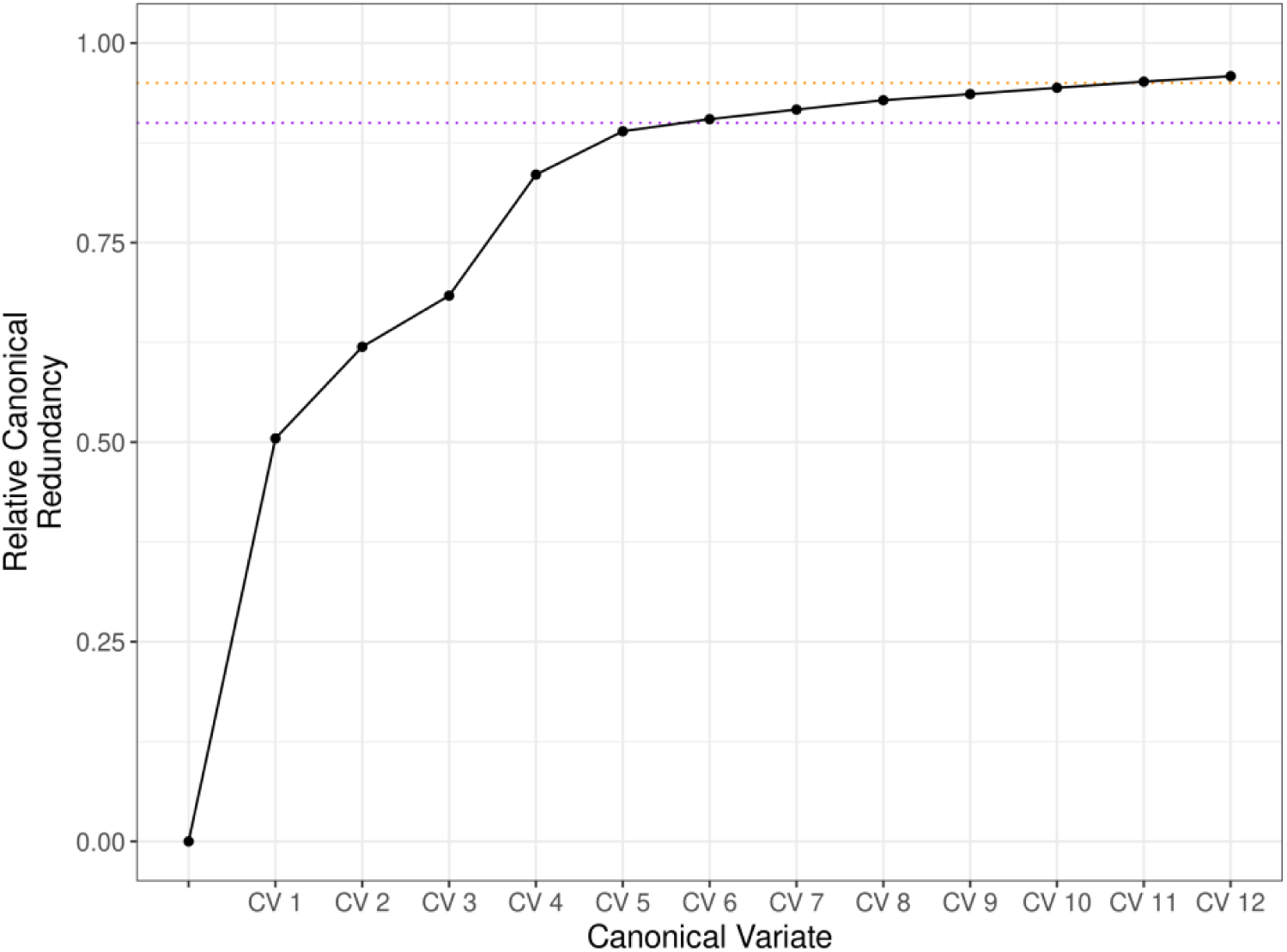
Cross-mate CCA scree plot based on alternative multiple imputation scheme. Vertical axis reflects sex-averaged cumulative relative canonical redundancy. Again, twelve linearly independent canonical variates are required to account for 95% of the variance explicable in one mate’s phenotype by linear combinations of their partner’s phenotypes. Here, missing values were imputed separately in males in females across five multiple imputations with chained equations using Random Forests predictions. Values reflect median redundancies across 5 × 5 imputed datasets.

**Figure S3.**
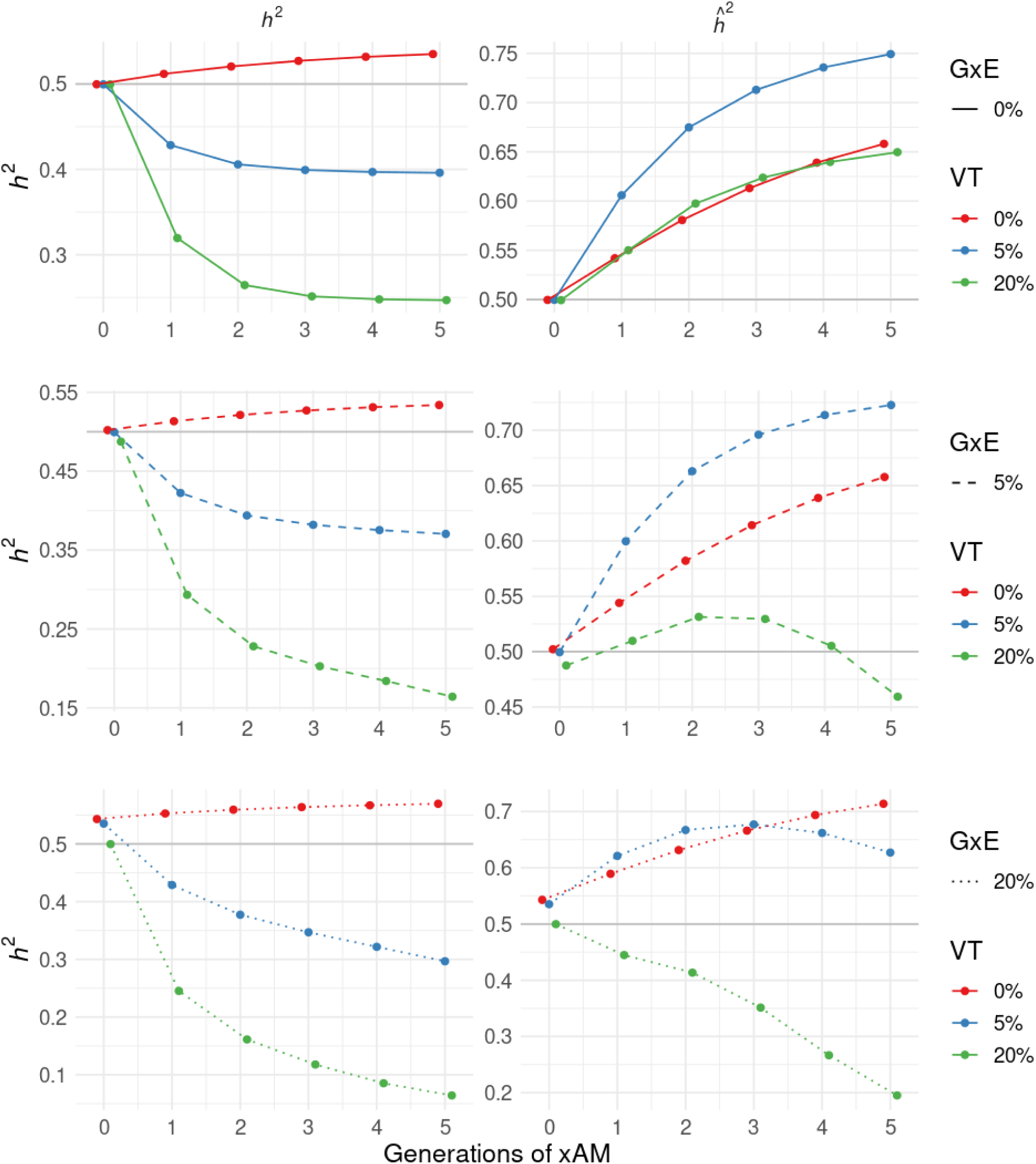
True and estimated heritability across successive generations of 5-variate xAM averaged over five phenotypes with orthogonal genetic effects. Architectures include gene-environment interactions (G×E; varying across rows), vertical transmission of environment from parent to offspring (VT), or both, in addition to cross-trait assortative mating (xAM). When present, G×E and VT account for 5% or 20% of the total phenotypic variance at generation zero.

**Figure S4.**
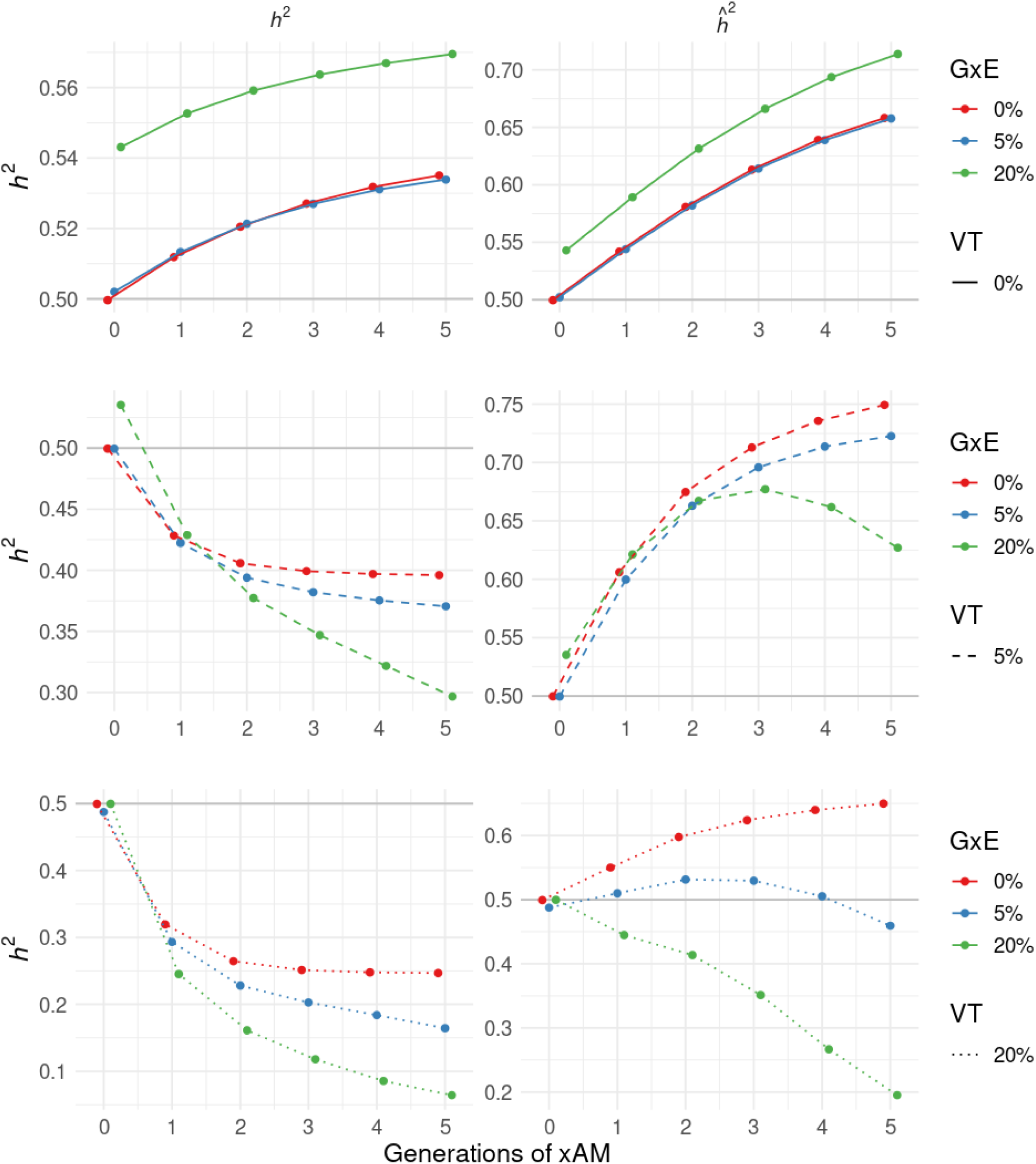
Alternative presentation of true and estimated heritability across successive generations of 5-variate xAM averaged over five phenotypes with orthogonal genetic effects. Architectures include gene-environment interactions (G×E), vertical transmission of environment from parent to offspring (VT; varying across rows), or both, in addition to cross-trait assortative mating (xAM). When present, G×E and VT account for 5% or 20% of the total phenotypic variance at generation zero.

**Figure S5.**
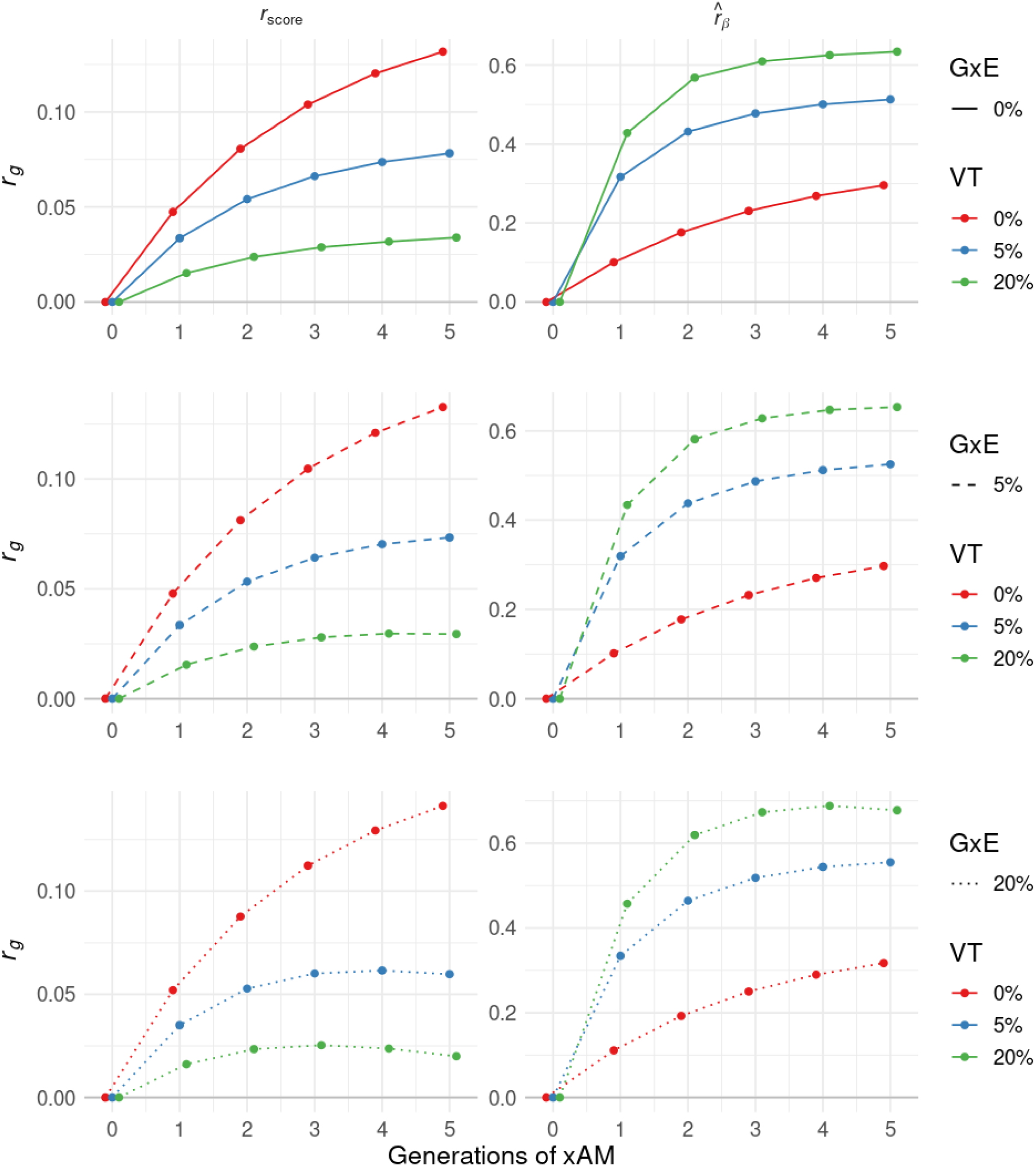
Alternative presentation of true polygenic score correlation and estimated effect correlation across successive generations of 5-variate xAM averaged over five phenotypes with orthogonal genetic effects. Architectures include gene-environment interactions (G×E), vertical transmission of environment from parent to offspring (VT; varying across rows), or both, in addition to cross-trait assortative mating (xAM). When present, G×E and VT account for 5% or 20% of the total phenotypic variance at generation zero.

**Figure S6.**
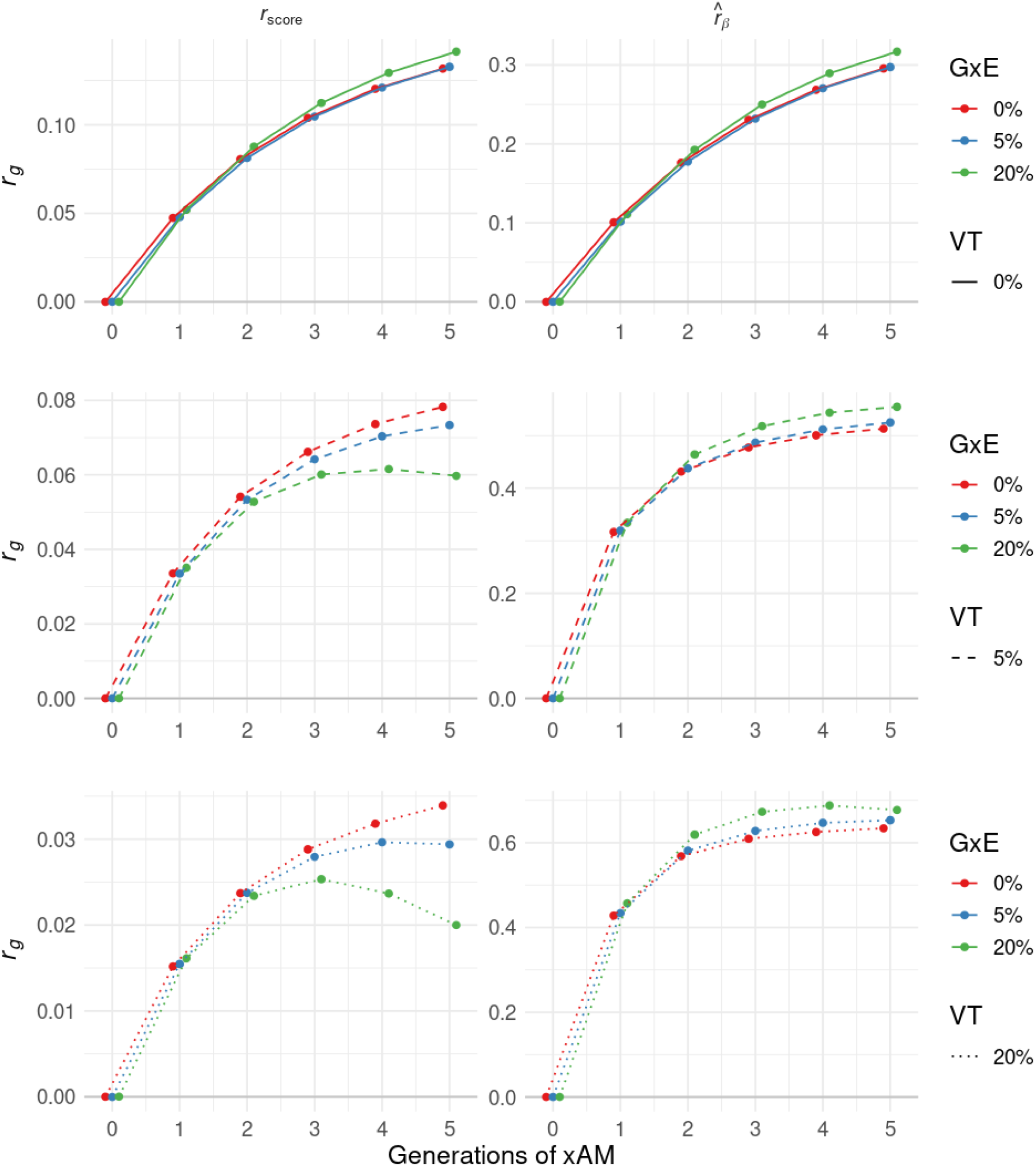
True polygenic index correlation and estimated effect correlation across successive generations of 5-variate xAM averaged over five phenotypes with orthogonal genetic effects. Architectures include gene-environment interactions (G×E), vertical transmission of environment from parent to offspring (VT; varying across rows), or both, in addition to cross-trait assortative mating (xAM). When present, G×E and VT account for 5% or 20% of the total phenotypic variance at generation zero.

**Figure S7.**
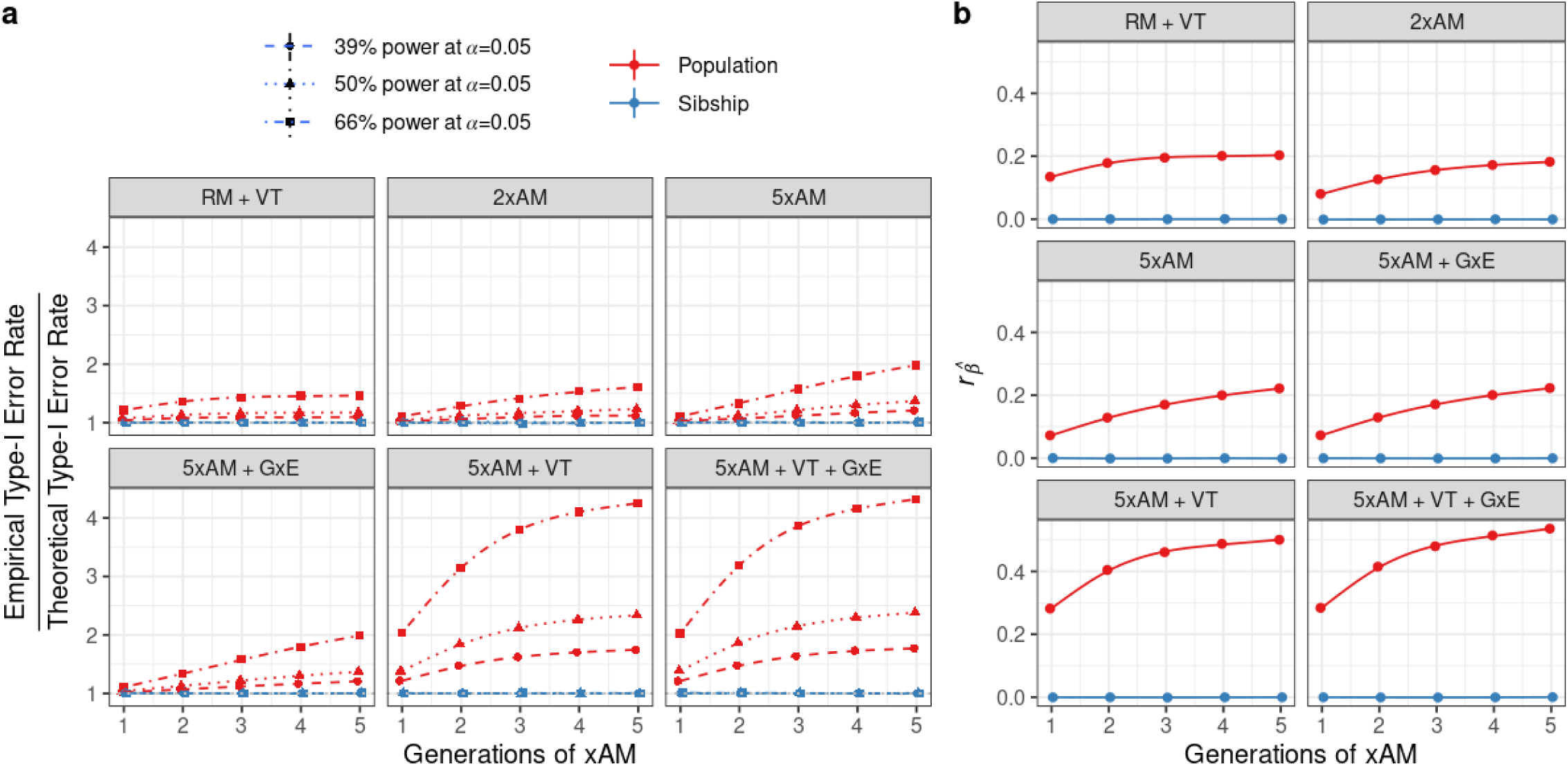
Population versus within-sibship GWAS. **(a)** Relative inflation of empirical to nominal type-I error rates at nominal significance for population-based and within-sibship GWAS with varying power (modulated here by the ratio of sample size to causal variants n/m) across successive generations. Simulations reflect random versus assortative mating (RM versus xAM) with or without gene-environment interactions (G×E) and/or vertical transmission (VT), all in the absence of pleiotropy. When present, G×E and VT account for 5% of the total phenotypic variance at generation zero. **(b)** Cross-phenotype correlations between population-based versus within sibship GWAS slope estimates over successive generations across simulation conditions.

**Figure S8.**
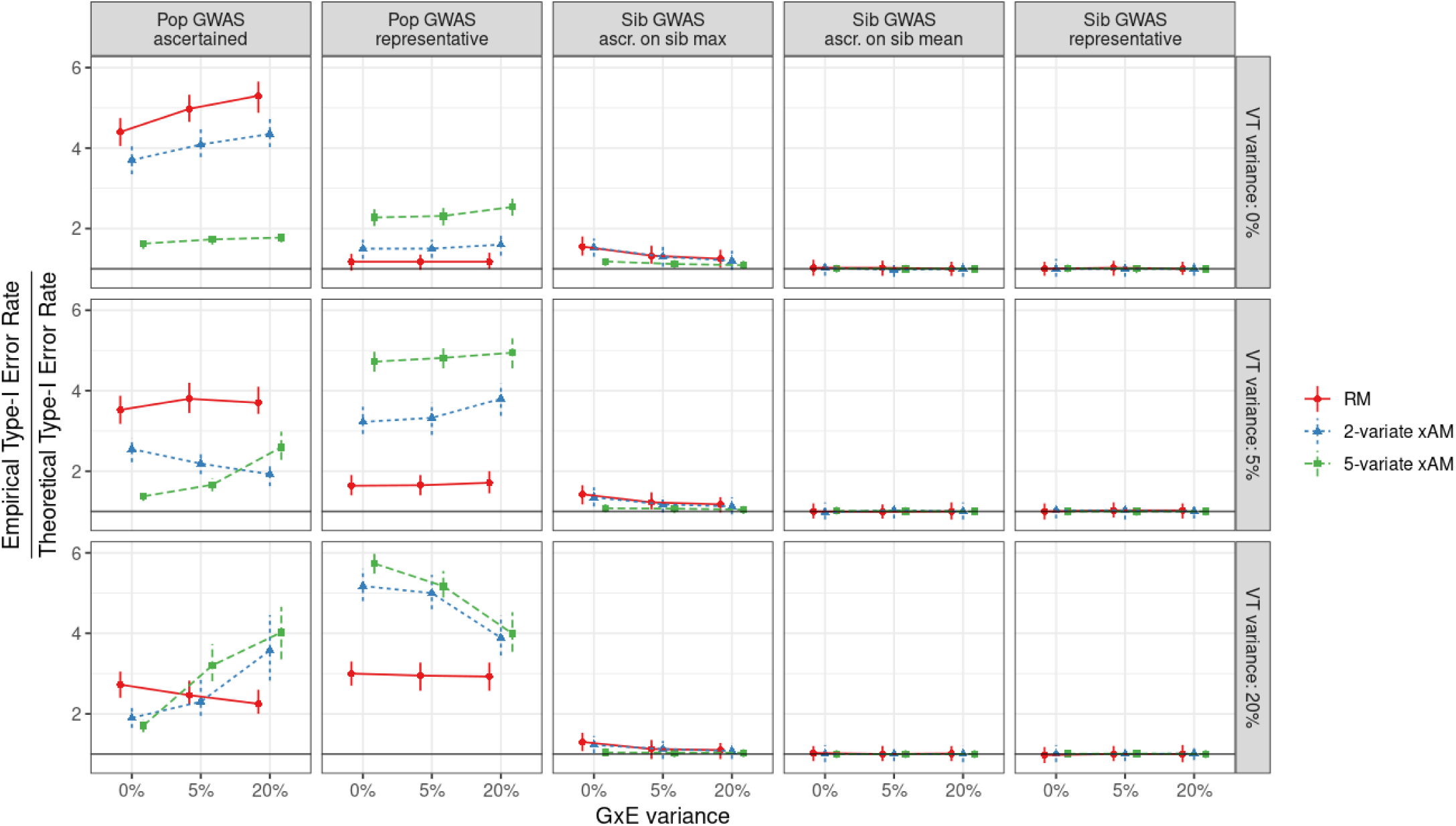
Relative inflation of empirical to nominal type-I error rates at nominal significance for within-sibship and population-based GWAS in representative samples versus ascertained samples where higher values are preferentially included. Simulations reflect varying mating regimes (random [RM] versus 2-and 5-variate cross-trait assortative mating [xAM] with exchangeable cross-mate cross-trait correlations of 0.2) across architectures with or without gene-environment interactions (G×E) and/or vertical transmission (VT), all in the absence of pleiotropy. When present, G×E and VT account for 5% or 20% of the total phenotypic variance at generation zero. In the case representative sampling, the within-sibship design mitigates the false positive rate inflation seen in population-based GWAS. Ascertainment is with respect to the sum of the 2 or 5 phenotypes (observations below the median value are discarded). Whereas ascertaining on sibling mean (i.e., the cross-sibling mean of these values) maintains this pattern, ascertaining on sibling maxima induces inflation and the extent of this inflation varies as a complex function of mating regime and the strength of G×E and VT effects.

**Figure S9.**
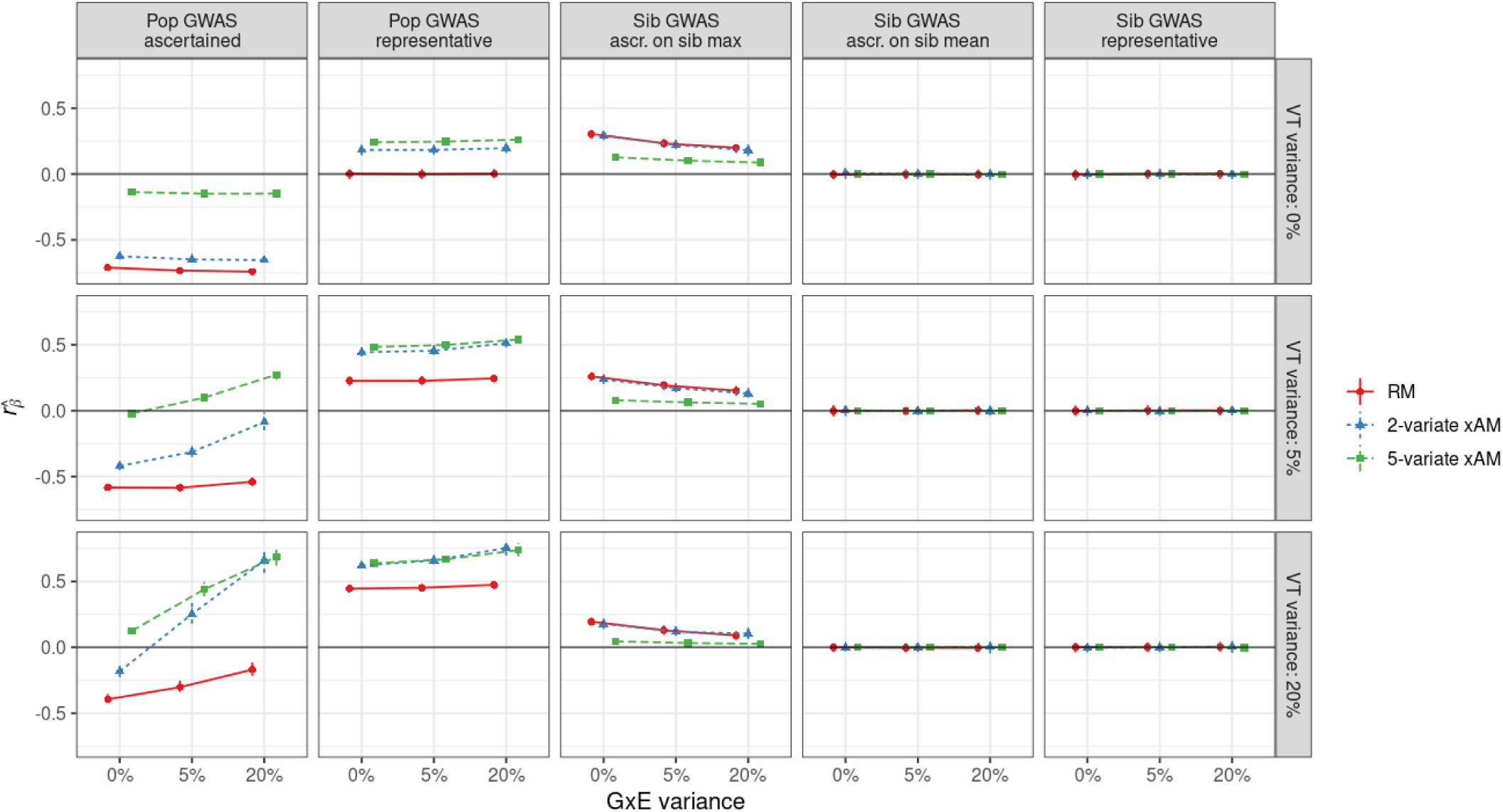
Correlation between within-sibship and population GWAS slope estimates in representative samples versus ascertained samples where higher values are preferentially included. Simulations reflect varying mating regimes (random [RM] versus 2-and 5-variate cross-trait assortative mating [xAM] with exchangeable cross-mate cross-trait correlations of 0.2) across architectures with or without gene-environment interactions (G×E) and/or vertical transmission (VT), all in the absence of pleiotropy (i.e., the true correlation slope correlation is zero). When present, G×E and VT account for 5% or 20% of the total phenotypic variance at generation zero. In the case of representative sampling, the within-sibship design mitigates the bias observed in population-based GWAS. Ascertainment is with respect to the sum of the 2 or 5 phenotypes (observations below the median value are discarded). Whereas ascertaining on sibling mean (i.e., the cross-sibling mean of these values) maintains this pattern, ascertaining on sibling maxima induces inflation and the extent of this inflation varies as a complex function of mating regime and the strength of G×E and VT effects.

**Figure S10.**
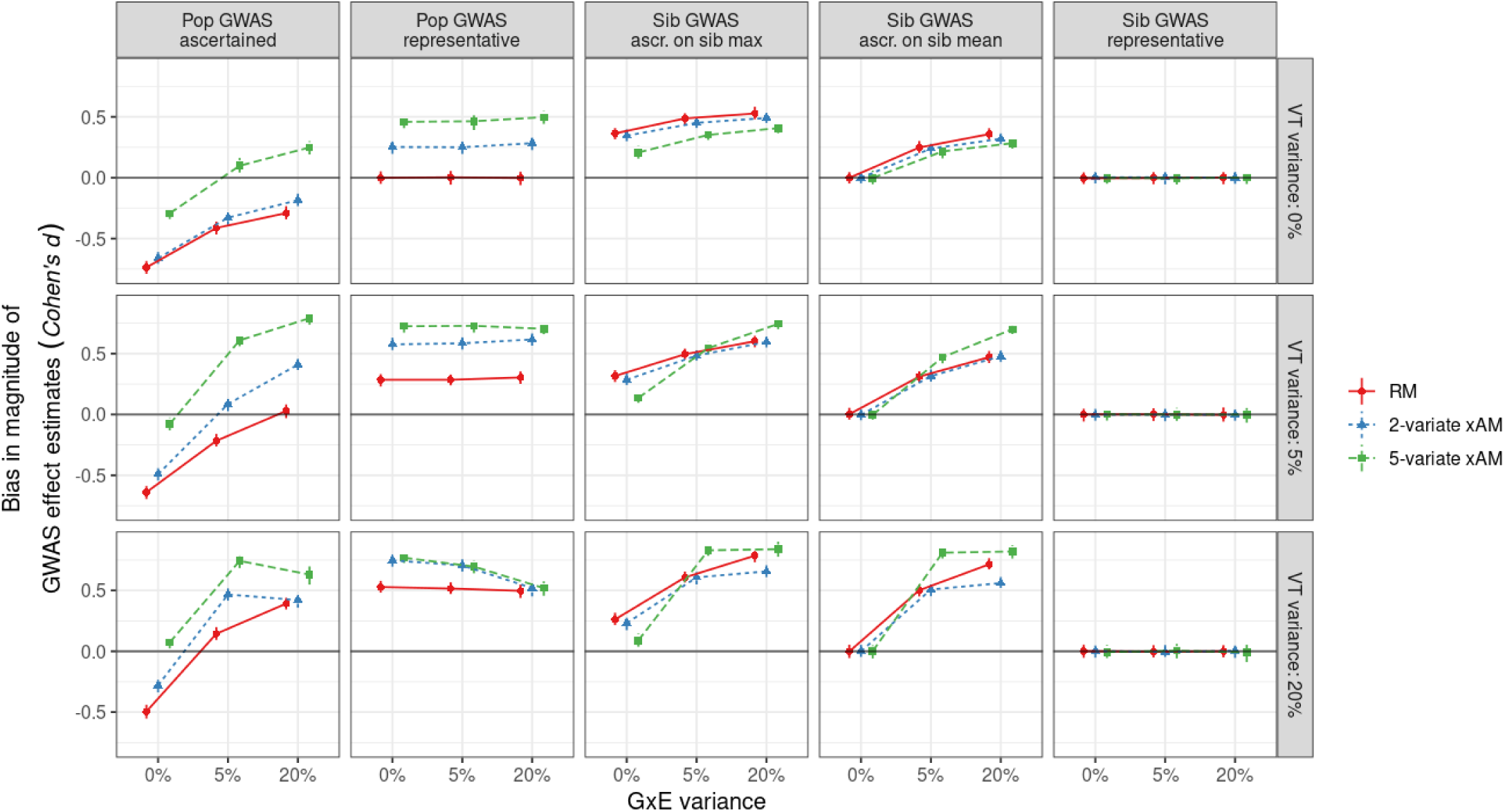
Bias in magnitude of estimated GWAS slopes for within-sibship and population-based GWAS in representative samples versus ascertained samples where higher values are preferentially included (observations below the median value are discarded). Bias is quantified using Cohen’s d. Simulations reflect varying mating regimes (random [RM] versus 2-and 5-variate cross-trait assortative mating [xAM] with exchangeable cross-mate cross-trait correlations of 0.2) across architectures with or without gene-environment interactions (G×E) and/or vertical transmission (VT), all in the absence of pleiotropy. When present, G×E and VT account for 5% or 20% of the total phenotypic variance at generation zero. In contrast to false positive rate inflation (Figure S8) and effect estimate correlations (Figure S9), both mean- and max-based ascertainment can induce artifacts in within-sibship GWAS.

**Figure S11.**
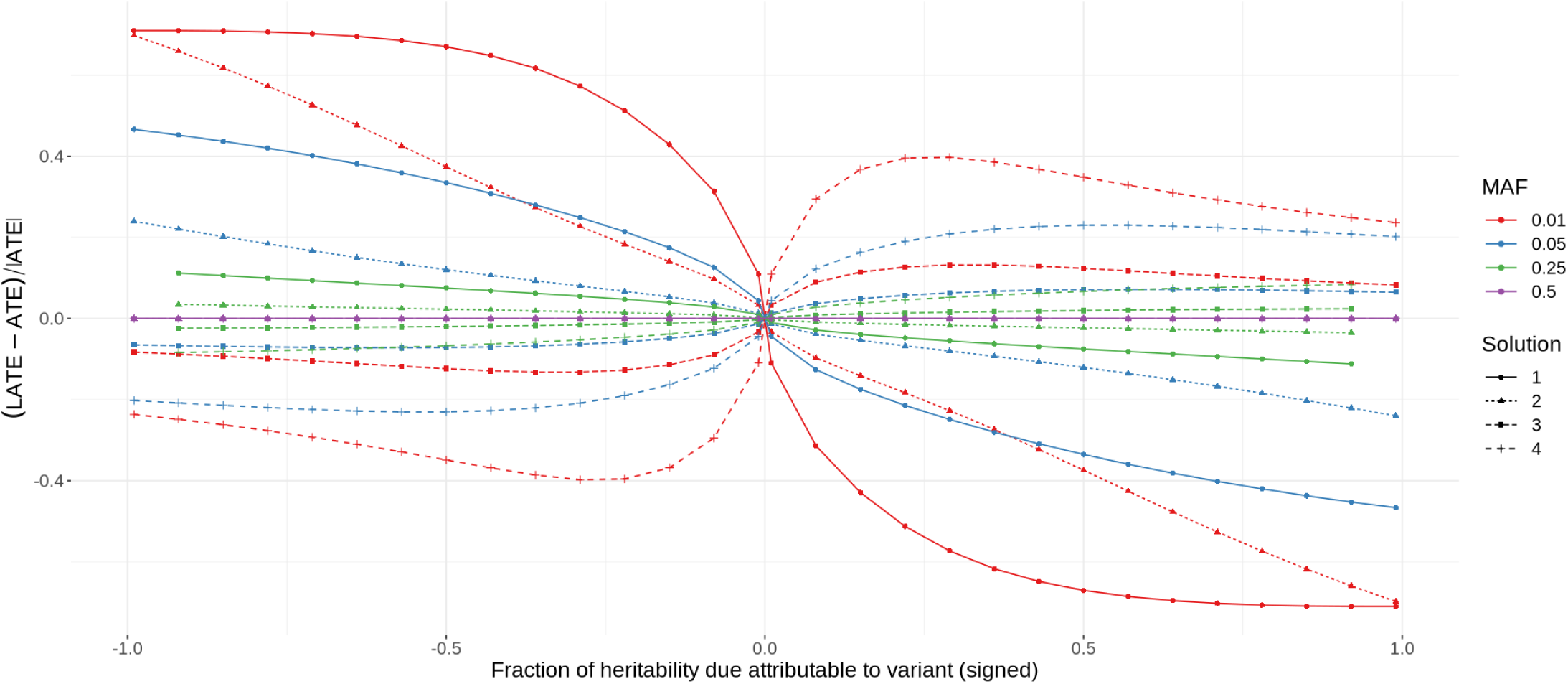
Relative difference between the local average treatment effect (LATE) for children of heterozygous parents and average treatment effect (ATE) of a single variant as a function of the fraction of heritability it accounts for, signed according to effect direction, for varying minor allele frequencies (MAF). Whereas the ATE reflects the average causal effect in the entire population, the LATE reflects that among individuals with one or more heterozygous parents at a given locus. Sibling-difference GWAS estimates the LATE as doubly-homozygous parents produce offspring with identical genotypes at the corresponding locus. In all cases, the total additive genetic, random noise, vertical transmission, and G×E variance components are respectively fixed to 0.5, 0.3, 0.1, and 0.1. For each MAF-effect combination, variance components alone do not uniquely determine a linear vertical transmission + G×E generative model, and instead correspond to four possible parametrizations, counting multiplicities. Points reflect numeric solutions obtained using a computer algebra system; missing values are an artifact of numerical instabilities at particular combinations of parameters.

**Figure S12.**
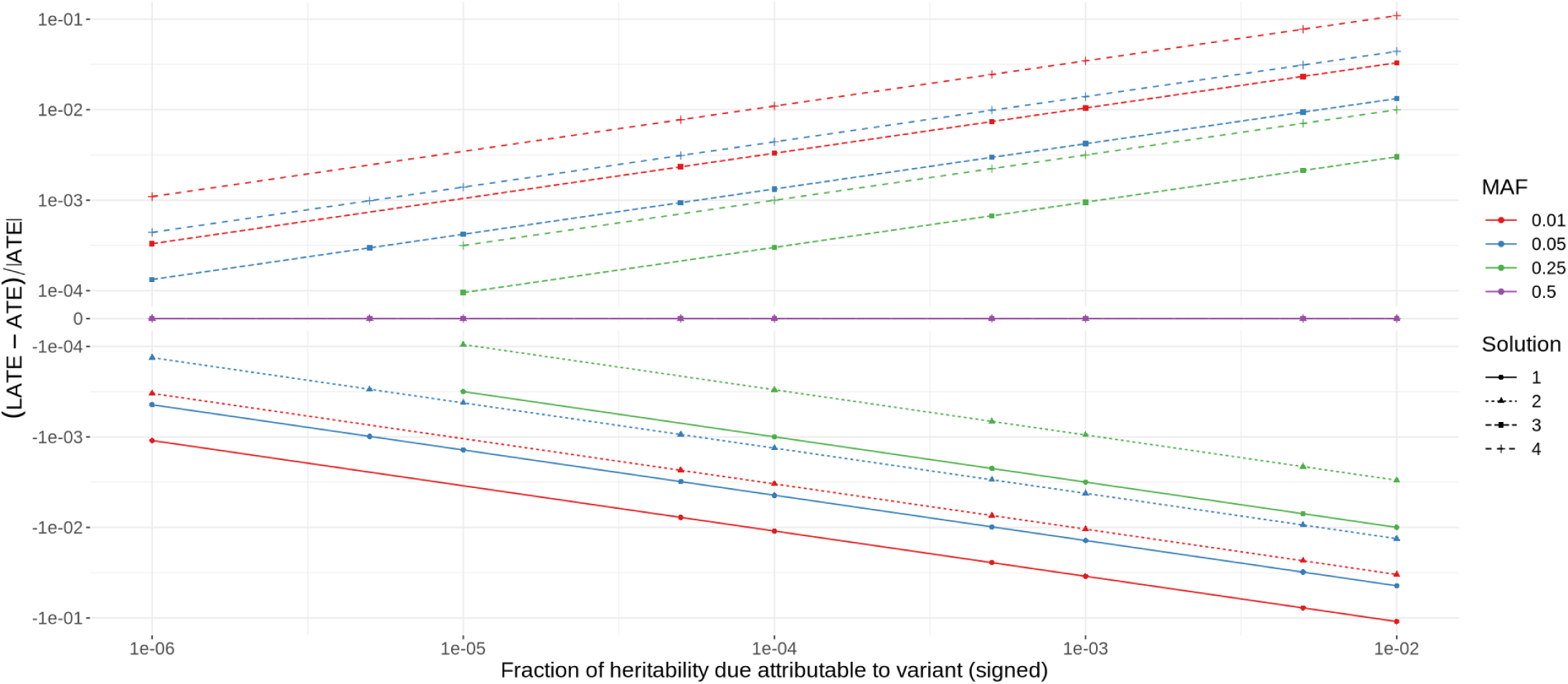
Relative difference between the local average treatment effect (LATE) for children of heterozygous parents and average treatment effect (ATE) of a single variant as a function of the fraction of heritability it accounts for, signed according to effect direction, for varying minor allele frequencies (MAF), on a pseudo-log^10^ scale. Whereas the ATE reflects the average causal effect in the entire population, the LATE reflect that among individuals with one or more heterozygous parents at a given locus. Sibling-difference GWAS estimates the LATE as doubly-homozygous parents produce offspring with identical genotypes at the corresponding locus. In all cases, the total additive genetic, random noise, vertical transmission, and G×E variance components are respectively fixed to 0.5, 0.3, 0.1, and 0.1. For each MAF-effect combination, variance components alone do not uniquely determine a linear vertical transmission + G×E generative model, and instead correspond to four possible parametrizations, counting multiplicities. Points reflect numeric solutions obtained using a computer algebra system; missing values are an artifact of numerical instabilities at particular combinations of parameters.

**Figure S13.**
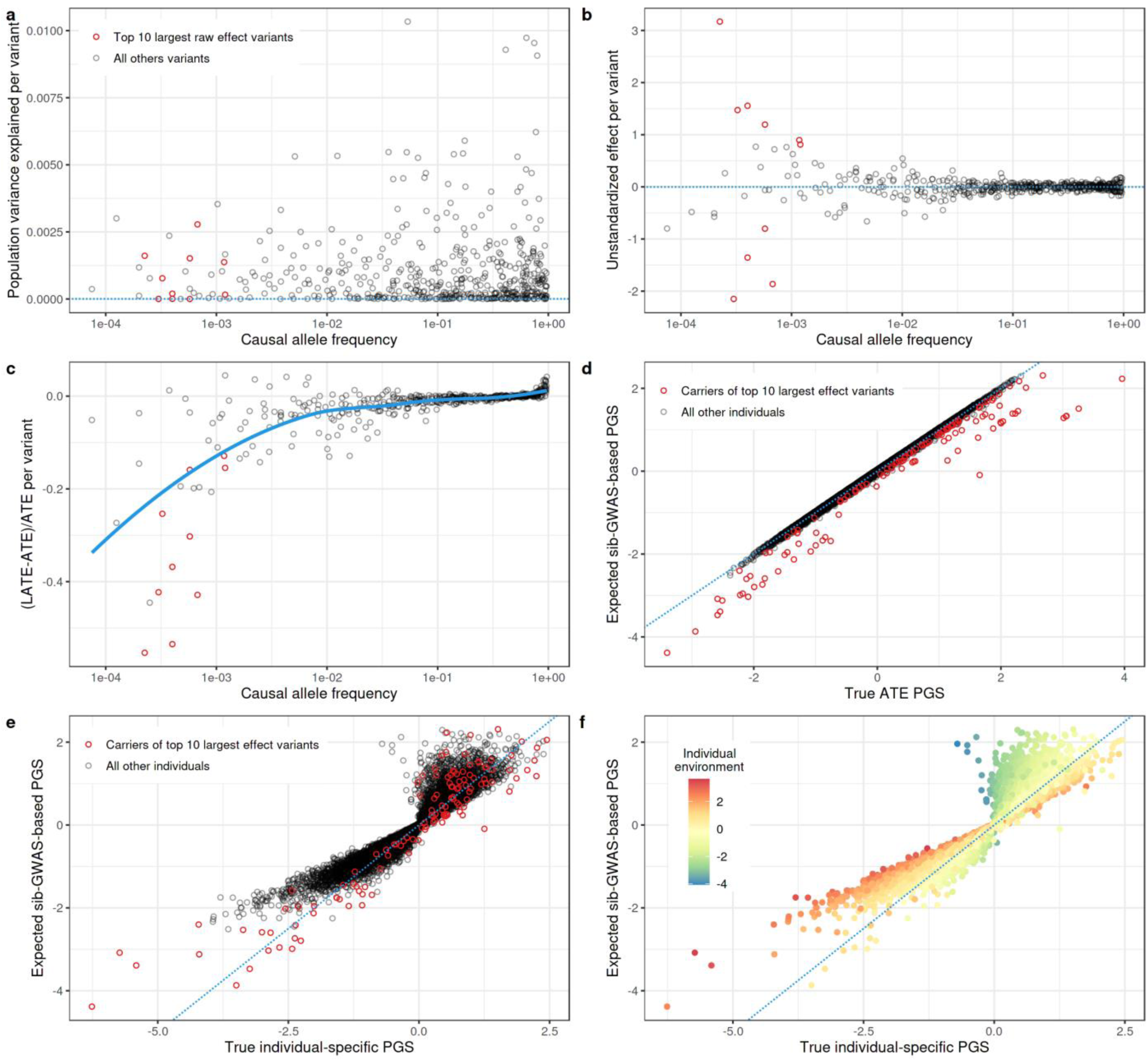
Vertical transmission and G×E induced bias of sibling-difference GWAS-derived PGI for carriers of large-effect variants when rare variants tend to have deleterious effects (see Methods for details of generative model). Despite (**a**) explaining little phenotypic variance in the population, (**b**) the top ten largest unstandardized effect variants (**c**) have substantially different sibling-GWAS estimated effects (the local average treatment effect [LATE] for individuals with one or more heterozygous parents) versus average treatment effects (ATE) in the population. As a result, (**d**) sibling-difference GWAS derived PGI for carriers of these rare large-effect variants yield substantially different values from PGI constructed using true ATE values. However, neither ATE- or LATE-based PGI fully characterize individual genetic liability in the presence of G×E (i.e., within individual-specific environmental contexts). Though not specific to family-based GWAS methods and **(e)** separate from the phenomenon inducing discrepancies among LATE-versus ATE-based PGI for carriers of large effect variants, **(f)** all additive PGI to account for individual environmental context and yield compromised estimates for individuals at environmental extremes.

**Figure S14.**
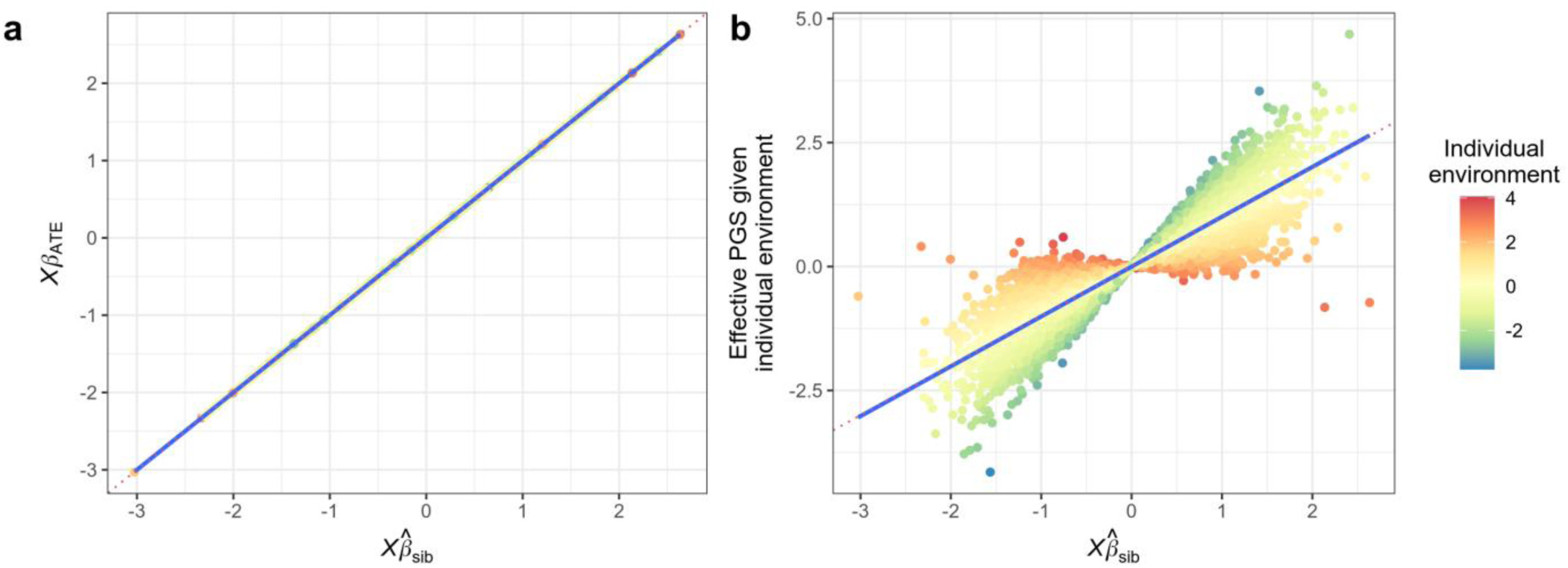
Individuals’ true average treatment effect (ATE) PGI (Xβ_ATE1_ ≔ E[Xβ|E = E̅]), which averages across environments, versus true individual-specific PGI (i.e., in the context of individual-specific environments), as a function of sibling-difference GWAS derived PGI, in the presence of G×E. In all cases, the total additive genetic, random noise, and G×E variance components are respectively fixed to 0.5, 0.4, and 0.1 (See Methods for generative model). In this setting (which notably lacks variants of large effect), despite the fact that individual effect estimates differ from true ATEs (Supplemental Figures S11-S12), these differences average out across the genome and **(a)** sibling-difference GWAS based PGI provide unbiased estimates of individuals’ ATE-based PGI. **(b)** On the other hand, unbiased ATE-based PGI will still yield misleading predictions under G×E for individuals in extreme environments (here colored in terms of standard deviations from the population mean). This phenomenon is not unique to sibling-difference GWAS and will affect all polygenic predictors that do not account for individual environment.

**Figure S15.**
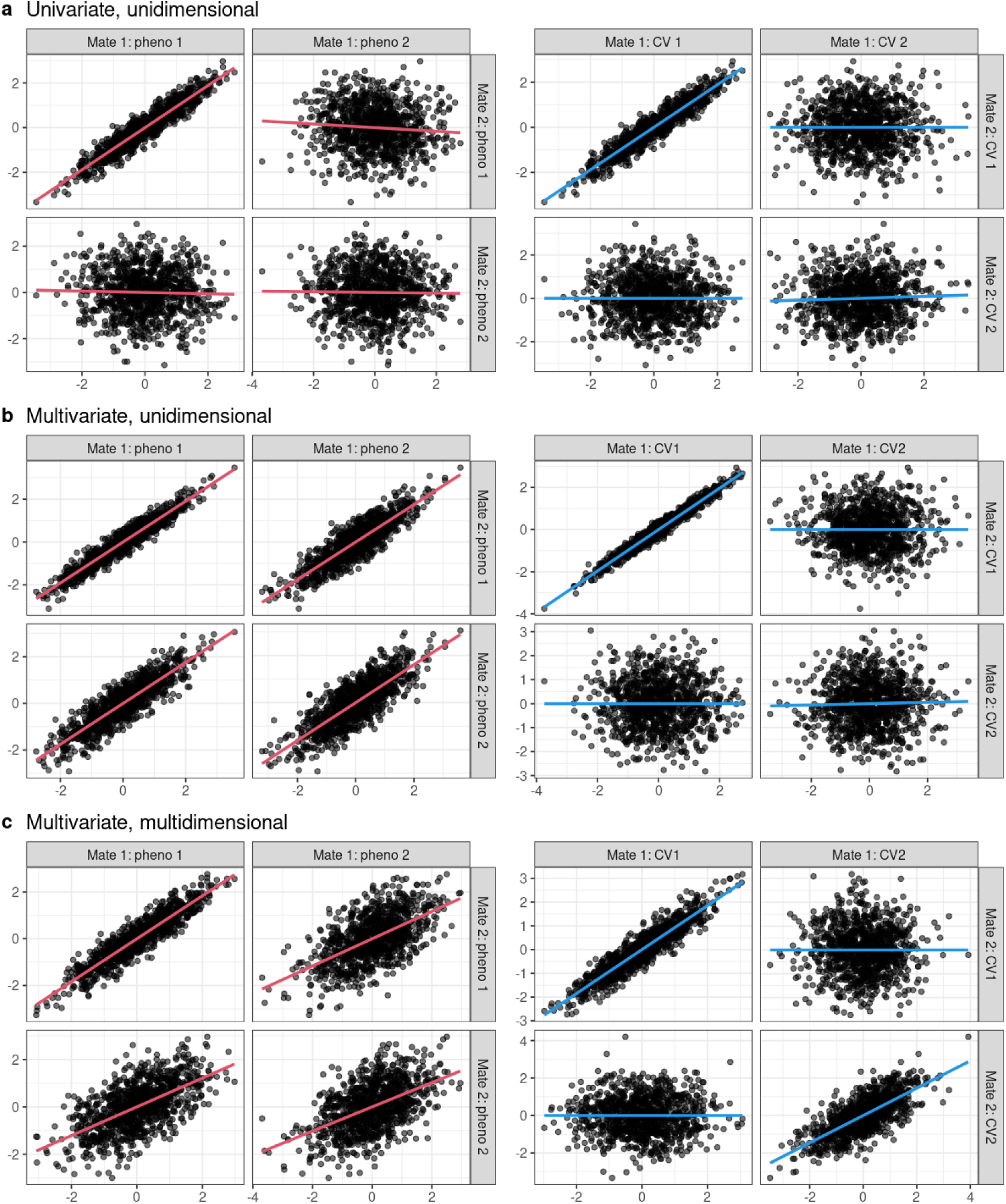
Multivariate versus multidimensional linear mating regimes. Each pane features cross-mate pair plots synthetic phenotypes (left) and corresponding canonical variates (CVs, right) with ordinary least squares best fit lines. **(a)** A univariate, unidimensional mating regime: mates are similar only on phenotype 1, and the first CV captures this covariance. **(b)** A multivariate, unidimensional mating regime: though mates are similar with respect to multiple phenotypes, this similarity is captured by a single CV. **(b)** A multivariate, multidimensional mating regime: mates are similar with respect to multiple phenotypes and multiple CVs are required to account for their similarity.

**Figure S16.**
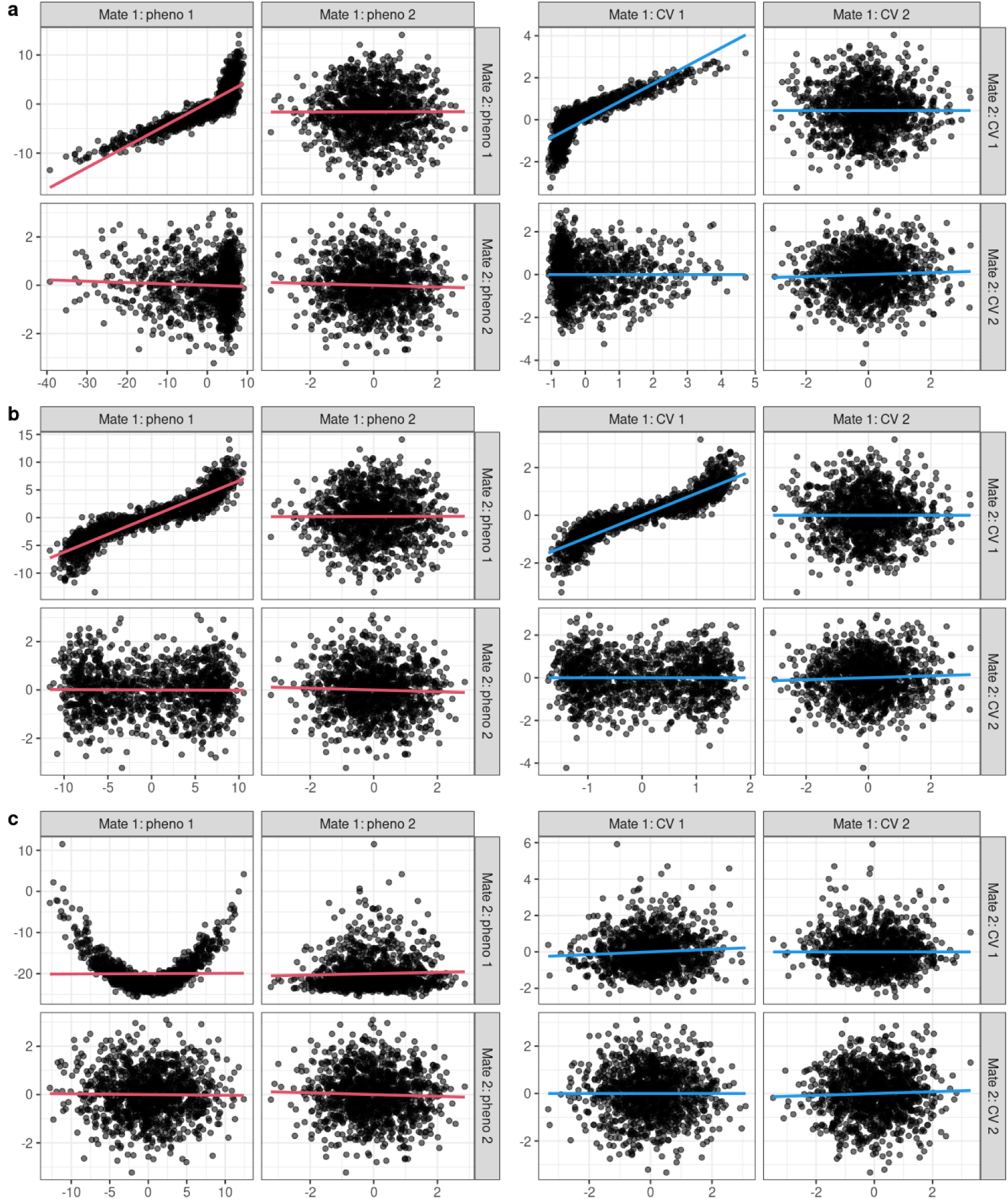
Canonical correlation analysis of univariate nonlinear mating regimes. Each pane features cross-mate pair plots synthetic phenotypes (left) and corresponding canonical variates (CVs, right) with ordinary least squares best fit lines. One mate’s phenotype is a **(a)** piece-wise linear, **(b)** inverse tangent, or **(c)** quadratic transformation of the other’s. These simple transformations fail to increase the apparent dimensionality: the monotonic transformations (a/b) are largely captured by a single CV and the non-monotonic transformation is not captured at all.

